# Resource-diversity relationships in bacterial communities reflect the network structure of microbial metabolism

**DOI:** 10.1101/2020.09.12.294660

**Authors:** Martina Dal Bello, Hyunseok Lee, Akshit Goyal, Jeff Gore

## Abstract

The relationship between the number of available nutrients and community diversity is a central question for ecological research that remains unanswered. Here, we studied the assembly of hundreds of soil-derived microbial communities on a wide range of well-defined resource environments, from single carbon sources to combinations of up to 16. We found that, while single resources supported multispecies communities varying from 8 to 40 taxa, mean community richness increased only one-by-one with additional resources. Cross-feeding could reconcile these seemingly contrasting observations, with the metabolic network seeded by the supplied resources explaining the changes in richness due to both the identity and the number of resources, as well as the distribution of taxa across different communities. By using a consumer-resource model incorporating the inferred cross-feeding network, we provide further theoretical support to our observations and a framework to link the type and number of environmental resources to microbial community diversity.

---

Uncovering the determinants of community diversity is central in ecology^1–3^ and microbiome research^4^, posing unique challenges to microbial ecologists. Indeed, microbes are the most abundant form of life on our planet^5^, the most ancient and the most phylogenetically diverse^6^. Surveys of a variety of ecosystems, from oceans^7^ to the human body^8^, have revealed that thousands of different taxa can stably coexist within the same community. Importantly, microbial communities drive the bulk of global nutrient cycling^9^, sustain human health^10^ and modulate the response of the biosphere to climate change^11^. Hence, deepening the knowledge of the drivers of microbial community diversity is pivotal to understand the functioning of Earth’s ecosystems.

Several mechanisms contribute to the diversity of microbial communities, including the spatial and temporal structure of the environment^12^, dispersal and bacterial motility^13^, warfare^14,15^, and resource-mediated competition and cooperation^16–18^. With respect to resources, ecological theory has mostly focused on the effect of the number of available resources on community diversity rather than their identity^19^. In particular, according to the principle of competitive exclusion, the number of stably coexisting species is predicted to be bounded by the number of available resources^20–22^. Despite the wealth of theoretical work on how resources can affect microbial community diversity, empirical tests of resource-diversity relationships have been limited, having been explored either in 2-3 species assemblages^17^ or in enriched cultures grown on 1-2 resources^18,23–25^. Systematic experiments encompassing a range of resource combinations are still lacking.

While an empirical test of the relationship between the number of available nutrients and community diversity remained elusive, bottom-up experiments have implicated cross-feeding as a major factor influencing the assembly of microbial communities, even in simple environments. Cross-feeding, whereby metabolic byproducts of one taxa become resources for others^26^, can increase niche partitioning, ultimately allowing the coexistence of several taxa even when only a single source of carbon is provided^18,23,25,27^. There is also some evidence that the identity of the supplied resource dictates community composition, as microbial taxa display different resource preferences and patterns of metabolite excretion^23,28^. Nevertheless, the manner in which cross-feeding and niche partitioning systematically change with the identity and the number of supplied resources is still unclear. This lack of knowledge impairs our ability to link variations in resource availability with shifts in microbial community diversity.

Here, we used a high-throughput experimental protocol and 16S amplicon sequencing to explore the relationship between microbial community diversity and resource availability in experimental microcosms. By growing soil-derived communities in media containing different combinations of carbon sources (from single resources up to 16), we discovered that community diversity was high in single resources but then increased only modestly with additional nutrients. These seemingly contrasting observations reflected the structure of the metabolic network seeded by the supplied resources. Cross-fed byproducts predicted to originate from each resource via microbial metabolism were coupled to the richness and composition of single resource communities. Additionally, a consumer-resource model incorporating the inferred metabolic network recapitulated the linear increase of community diversity with additional resources.

## Results

In order to illuminate how the availability of resources, namely their number and identity, shape the richness of microbiomes, we assayed the assembly of soil-derived bacterial communities in laboratory microcosms^16,18^. We started by inoculating a rich microbial suspension obtained from a soil sample (Fig. S1) into 75 resource environments, each containing minimal media supplemented with different combinations of carbon sources, ranging from one to 16 (Fig. 1a, S2, Table S1). The 16 carbon sources represented a broad range of common soil compounds (e.g., mannose, xylose, cellulose and hydroxyproline), encompassing both glycolytic (e.g., simple and complex sugars) and gluconeogenic substrata (e.g., organic acids). We adopted a daily-dilution protocol, whereby at the end of each 24-hour growth cycle the bacterial cultures were diluted 1/30x into fresh media. We observed that the majority of microcosms reached stability after 3 days from the inoculum (Fig. S3). We continued the experiment until day 7 and measured the final richness as the number of ASVs (amplicon sequence variants) observed within each community (Fig. S4).

**Figure 1.**
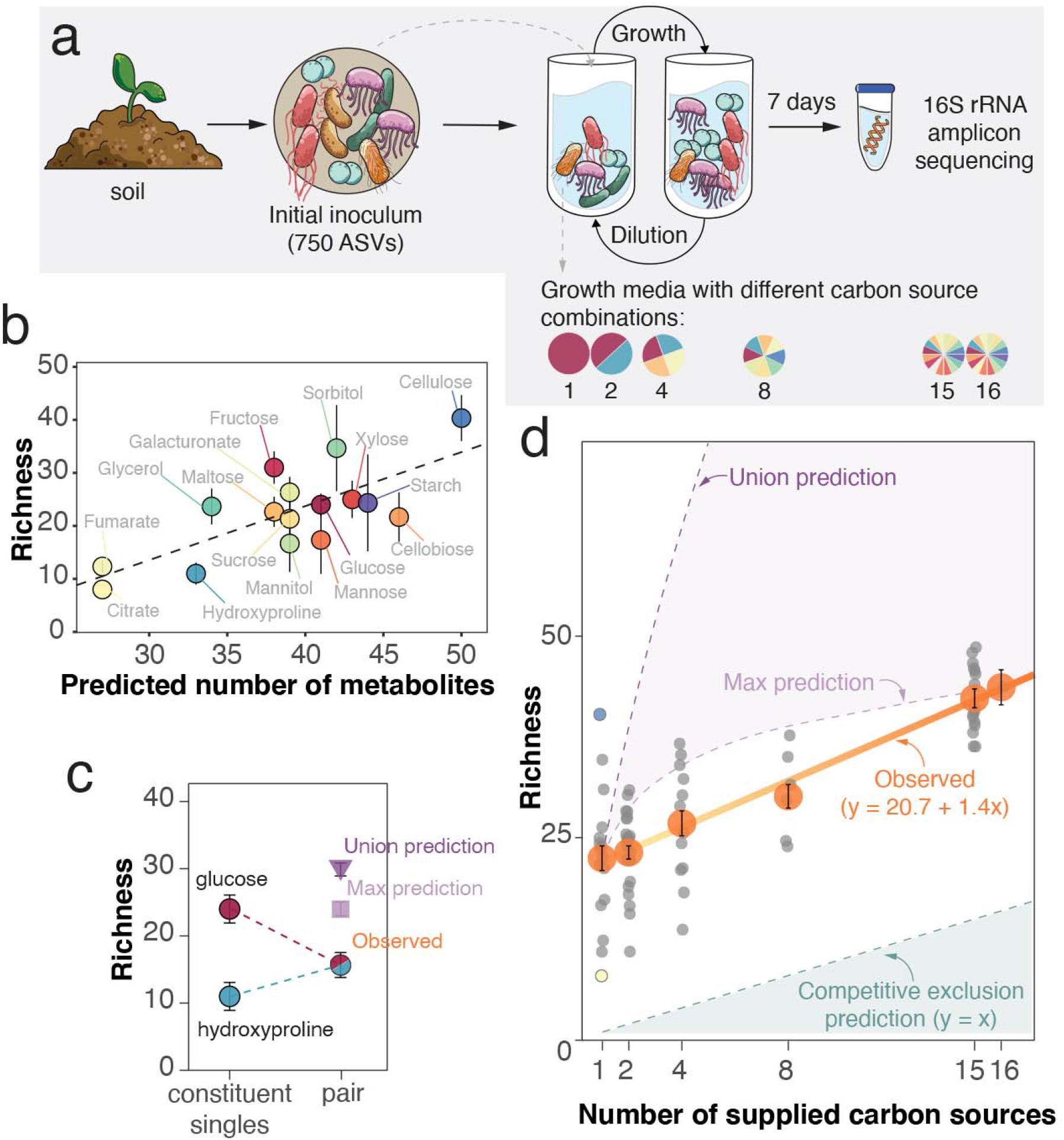
Microbial community diversity increases slowly with the number of resources despite individual resources supporting complex multispecies communities. **a**. Layout of the experiment. We inoculated a rich microbial suspension obtained from a soil sample into 75 growth media, each supplemented with a different combination of carbon sources, from single compounds to a mix of 16, while keeping the total carbon concentration the same (0.1% w/v). Bacterial cultures were grown for 7 days under a regime of daily dilution and their composition assessed at the single nucleotide resolution using 16S rRNA amplicon sequencing. Community diversity is measured as richness, i.e., number of AVSs. **b**. Richness of microbial communities supported by single carbon sources correlates with the number of metabolites predicted to be generated from metabolic reactions mapped in the KEGG database (Pearson’s correlation coefficient r=0.75 [95% CI: 0.4-0.91], p<0.001). Colored dots indicate, for each carbon source, the number of ASVs (mean ± SEM, N = 3). **c**. A representative example of how observed richness in constituent single resources (mean ± SEM, N = 3) compares to the observed richness in two-resource communities (mean ± SEM, N = 3) and predictions calculated as the union (sum without overlapping ASVs, dark violet) or the maximum (light violet) of the richness in constituent singles (mean ± SEM, N from permutations = 9). **d**. Observed average richness (orange dots, mean ± SEM, N = 16 for single-resource, 24 for two-resource, 12 for four-resource, six for eight-resource, 16 for 15-resource and 1 for 16-resource combinations) as a linear function of the number of supplied carbon sources (solid orange line). Grey jittered dots indicate the average richness for each unique combination of resources (mean ± SEM, N = 3). Intercept = 20.7± 0.8, slope 1.4 ± 0.1 (p <0.001). In single resources, the blue and yellow dots correspond to the highest and lowest average richness, measured in cellulose and citrate, respectively. The predicted trajectory of richness based on the competitive exclusion principle (dashed dark green line), the union (dashed dark violet line) and maximum (dashed light violet line) estimates, as described for panel b, are shown for comparison.

### Individually-supplied resources support complex multi-species communities

Consistent with recent experimental studies^18,24,29^, single-resource communities were remarkably rich (mean richness = 23 ± 2 ASVs, Fig. 1b) and taxonomically diverse (Fig. S5). This is in contrast with competitive exclusion predicting that the number of species cannot exceed the number of resources^20,30^—which, in single carbon sources, would result in no more than one species surviving. Interestingly, the variability in richness among different resources was also high—with the average number of ASVs ranging from 8 in citrate to 40 ± 4 in cellulose (Fig. S6, Fig. 1b)—and larger than the variability among replicates of the same carbon source (ANOVA test, *F*_*resource*_ = 3.4339, *p < 0*.*01*). Richness in single carbon sources therefore depended on resource identity. Community richness did not correlate significantly with the molecular weight of the supplied resource (Fig. S7), but did correlate with the predicted number of metabolites which could be generated from the resource through intracellular biochemical reactions and secreted in the environment (Fig. 1b, see Methods for the details on the prediction of metabolites based on KEGG^31^ and MetaCyc^32^ databases). Notably, the lowest richness was observed for gluconeogenic substrata (∼10 for citrate, fumarate and hydroxyproline), which were connected to the central metabolic pathway via the TCA cycle, hence resulting in the smallest metabolite pools. Consistent with previous work, these results highlight the role of cross-feeding in supporting community diversity^33–35^. Moreover, they suggest that the extent of cross-feeding may determine how many species can coexist on single resources.

Having found large numbers of coexisting species in single resources, we expected that community diversity would increase rapidly if more resources were provided. As previously observed in marine bacteria^24,25^, community composition could potentially correspond to the sum of the assemblages observed on each nutrient supplied in the mixture. To provide an example, the expected richness of the community grown on glucose and hydroxyproline (Fig. 1c), each alone supporting on average 24 and 11 ASVs, would be ∼ 30 ASVs, i.e., the sum minus the number of shared ASVs (union). Alternatively, niche overlap between the taxa found in the single-resource media^36,37^ might bring the expected number of species down to the maximum richness observed in the constituent singles; in the case of glucose + hydroxyproline, 24 ASVs.

However, when we measured the richness of the communities grown in a media supplied with equal amounts of glucose and hydroxyproline, we found only ∼16 ASVs on average, which is significantly lower than both expectations (Fig. 1c). Yet, our observed richness was remarkably similar to the mean richness measured in the two constituent single resources (17.5 ASVs), a trend that was consistent across many two-resource communities (Fig. S8). Contradicting our expectations based on previous results supporting additivity, we found that community richness upon combining two carbon resources was approximately the average richness of constituent single resource environments.

### Community diversity increases linearly with the number of supplied resources

Next, we examined the full range of resource combinations included in the experiment. Again, the richness predicted from the union of constituent singles significantly overestimated the observed richness (Fig. 1d). The prediction based on the maximum of constituent singles gave an increase with negative curvature that was not detected in our experiment (Fig. 1d). We found a similar trend also when we estimated the number of metabolites generated from resource combinations, with the same approach we used for single carbon sources (Fig. S9). Instead, the observed average richness increased linearly with the number of supplied carbon sources, at the constant rate of one to two ASVs for each new added resource (Fig. 1d, slope = 1.4 ± 0.1). As a result, the richness of communities supported by 16 resources was roughly twice the average richness of single-resource communities. The linear relationship was robust to the exclusion of low-abundance ASVs—with the slope reduced to 1 ± 0.07 when ASVs with relative abundance below 0.1% were excluded (Fig. S10a)—and coarse-graining at the family level (Fig. S10b). In addition, as more resources were provided, communities became more even (see Methods and Fig. S10c, d, S11), without changes in total biomass (Fig. S12). Despite confirming that the number of supplied resources is an important driver of microbial diversity, the observed one-by-one relation between richness and resource number was difficult to reconcile with the large diversity found in single resources. Thus, we went back to the single-resource communities to gain a better understanding of our observations.

### Communities are composed of generalists and variable numbers of specialists

First, we measured the resource occupancy of the 275 ASVs observed in single resource media, i.e., how many single-resource media a given ASV was found in (Fig. 2a, Fig. S13). Based on resource occupancy, we considered habitat specialists the ASVs that were observed in less than 25% of single-resource media, and habitat generalists those that occupied more than 75% of single-resource media (Fig. 2a). The majority of ASVs (216 out of 275) were specialists, whereas very few of them (10) were generalists. Some ASVs (49) displayed an intermediate occupancy, being present in between four to twelve media. This is reminiscent of natural communities, in which few taxa are usually universally present across different habitats, while the majority is found only under specific environmental conditions^38–40^. Importantly, previous work has shown that variations in the proportion of generalist and specialists taxa within a community impact its dynamics^41–43^.

**Figure 2.**
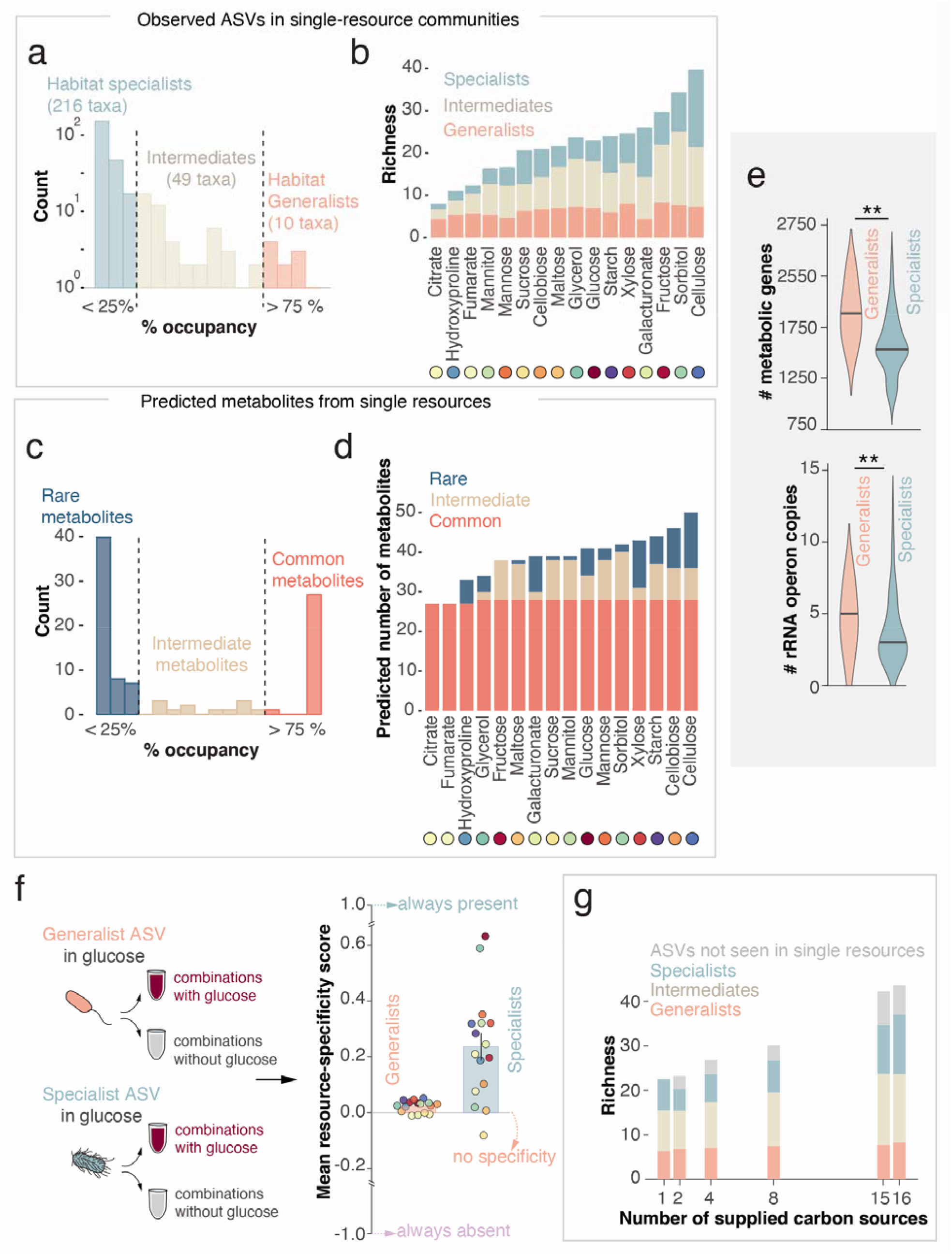
Experimental communities are composed of generalists and variable numbers of specialists, with the latter driving the increase in community diversity. **a**. The 275 ASVs found across all single-resource communities were classified in generalist, specialists and intermediates depending on their resource occupancy. The majority of ASVs exhibited a more specialized resource-utilization strategy. **b**. The richness in single resource-media is displayed highlighting the mean number of generalist (pink), intermediate (beige) and specialist (teal) ASVs (mean, N=3, error bars are omitted for clarity). **c**. The metabolites estimated to be produced starting from the supplied single resources through cell reactions can be classified in common, intermediate and rare, based on resource-occupancy as for ASVs. **d**. The total of metabolites estimated for each single resource is displayed highlighting the number of common (red), intermediate (light brown) and rare (blue) metabolites. **e**. Upper panel. The distribution of the number of metabolic genes retrieved for each ASV in single resources (see Methods) differs between generalists and specialists (p < 0.01, from Kolmogorov-Smirnov test). Lower panel. The distribution of rRNA operon copy numbers, calculated at the genus level, of generalist ASVs differs from that of specialist ASVs (p < 0.01, from Kolmogorov-Smirnov test). **f**. The specificity score is calculated, for each ASV found in a single resource (target resource), using the number of multi-resource media containing the target resource in which the ASV was found (X) and the number of media not containing the target resource in which the ASV is found (Y), as (X – Y)/(X + Y). It ranges from 1, indicating that the ASV is present only in a combination containing the resource, to −1, implying that the ASV is always absent when the resource is supplied. A score of 0 is indicative of an ASV showing no specificity for that particular resource. Bars indicate the mean specificity score ± SEM for generalists (pink) and specialists (teal) (N = 16). Colored dots indicate the mean score for each resource (SEM are omitted for clarity, N varies for each resource, see Methods). **g**. The mean number of generalist (pink), intermediate (beige) and specialist (teal) ASVs for media with the same number of resources is shown as stacked bars. The average number of ASVs that were not detected in single-resource communities but appeared in other combinations is indicated in grey. Error bars are omitted for clarity.

Next, we inspected the distribution of specialists and generalists within single resources. We found that species-poor communities, grown on gluconeogenic substrates like citrate, were dominated by generalist ASVs (often representing > 50% of observed taxa; Fig. 2b, S14a), while species-rich communities were enriched in specialists (see cellulose in Fig. 2b, S14b). We noticed that glycolytic substrates, which can produce a much larger metabolite pool before connecting with the central carbon metabolism, supported communities where more specialists coexisted with generalists. This suggested a link between the metabolite pool generated from each supplied resource and its ability to sustain both generalists and specialists in the same community.

Remarkably, just like ASVs, our predicted metabolic byproducts could also be broken up into two broad classes. Based on how many single resources could trigger their production, we could distinguish between common metabolites, present in association with the majority of single resources (like generalist taxa), and rarer metabolites, present only in association with one or few resources (like specialists) (Fig. 2c). Metabolites that were commonly produced constituted the *core* intermediates of the central metabolic pathway, including the TCA cycle and lower glycolysis. The rarely produced metabolites, instead, were the intermediates of *peripheral* branches of the central pathway. For example, if either citrate or fumarate were provided, we predicted that the central pathway proceeds in the gluconeogenic direction, generating only byproducts belonging to the core pool. In contrast, individually-supplied sugars were predicted to go through a series of reactions before entering the central pathway, ultimately generating both core and peripheral metabolites (Fig. 2d). It appeared that the number of peripheral metabolites varied with the position from which the resource entered the “metabolic map”. The parallelism in the distribution of ASVs and predicted metabolites reinforces the idea that community structure is coupled to the metabolite pool, and suggests a link between the resource occupancy and metabolic capability of taxa.

We next tested for systematic differences in the metabolic capabilities between generalist and specialist taxa in our experimental microcosms. Generalist ASVs belonged to the most abundant families, i.e., Pseudomonadaceae, Enterobacteriaceae and Micrococcaceae (Fig. S13) and differed metabolically from specialists, e.g., taxa from Cellvibrionaceae. In particular, generalists were estimated to harbor a larger number of metabolic genes (Fig. 2e upper panel, see Methods for details on the estimation of gene content) and more copies of the 16S rRNA operon compared to specialists (Fig. 2e lower panel, see Methods for details on the matching with the number of copies of the rRNA operon), indicative of faster max growth rates^44^. Both results are consistent with studies showing the hallmarks of a generalist life style: flexible metabolism^38,45^ (indicated by the number of metabolic genes) and capacity for fast growth (indicated by the 16S rRNA copy number)^46,47^. At the same time, several of the taxa classified as generalists are known to show distinct resource preferences when grown in isolation. For example, *Pseudomonads* species dominated in the communities sustained by organic acids, most likely because of their advantage over other taxa preferring sugars^48^, but were also present in all the media in which organic acids could have been generated as byproducts of the glycolytic metabolism of sugars^49^ (see Fig. S3). This might indicate that generalists were present in all the communities because the substrates that they utilize were always generated as byproducts of bacterial metabolism. Indeed, even habitat generalists show resource preferencies^50^, such as Pseudomonas spp., which consumes preferentially acetate and other organic acids^23^. In contrast, since many sugars and their intermediates could not be produced via gluconeogenic metabolism^51^, the survival of the taxa specializing on them was prevented unless those sugars were externally supplied. Together these observations are consistent with the idea of habitat generalists and specialists assembling in a community in relation to the available supplied and cross-fed metabolites.

The coupling between metabolite pool and community structure observed in single resources suggested that resource-ASVs associations would be maintained also in multi-resource environments. In particular, we expected that generalists would be present in all communities, while specialists would be mostly detected when the favorite substrate was provided or metabolically generated. To verify these expectations, we calculated a resource-specificity score. For each ASV present in a single resource (target resource), the resource specificity score was calculated as the difference between the number of multi-resource media containing the target resource in which the ASV was found and the number of media not containing the target resource in which the ASV was found, divided by the total number of media in which the ASV was found. The score ranged from 1, indicating that the ASV was present only when the target resource was provided in a combination, to −1, implying that, although the ASV was found in the single resource, it was always absent when that resource was supplied with others. A score of 0 indicated that an ASV showed no specificity for that resource (Fig. 2f). We found that specialists’ scores were on average positive across all resources (Fig. 2f, mean score = 0.24 ± 0.05), while generalists’ scores were on average nearly zero (0.02 ± 0.01). Together, these findings highlight that (specialist) taxa tend to show resource-specific associations, and that single resource-ASV associations are maintained even in multi-resource environments.

To verify how resource-ASV associations impacted the resource-diversity relationship, we then calculated the average number of specialists, generalists and intermediates (as defined based on single resource occupancy) for each combination of carbon sources. We found that going from 1 to 16 resources, communities went from containing a balanced mixture of generalists and specialists to being dominated by more specialized ASVs (both specialists and intermediates, Fig. 2g). Overall, these results point to the consistent coexistence in our experimental microcosms of distinct groups of bacteria, with more specialized taxa progressively favored by the supply of additional resources. While this was in line with the expectation that specialists of each resource should be favored by the higher chances to introduce a glycolytic compound as more resources were added, it is important to note that, at the same time, several specialist ASVs were lost and few new ASVs were introduced, especially going from one to two carbon sources (grey bars in Fig. 2f, S15, these ASVs remained unclassified).

In summary, our experimental results revealed that 1) single resources were able to sustain multispecies communities, 2) going from one to two resources, community richness did not significantly increase; 3) overall, the resource-diversity relationship was linear and only modestly increasing; 4) all experimental communities were composed of both habitat generalists and specialists and their ratio changed with the number of supplied resources. We also show that the structure of the metabolite pool, which is the result of the ensemble of metabolic reactions fueled by the supplied and cross-fed resource(s), is the most likely driver of the observed manifestations of the resource-diversity relationship. We next asked: can we recapitulate some of the primary features of our experimental results by incorporating the metabolic network in a resource-explicit modelling framework?

### A resource-explicit model incorporating a realistic metabolic network reproduces our experimental results

We hypothesized that the metabolic network seeded by the supplied carbon sources could explain the observed resource-diversity relationship. To test this, we implemented the well-known MacArthur consumer-resource model with cross-feeding^19,21,35^. In contrast to other implementations which used an abstract, randomly-generated cross-feeding network^18,35^, we used a realistic network inferred using KEGG and MetaCyc databases (Fig. 3a, see Methods for details). Note that we used the same network to estimate the possible number of metabolic byproducts generated from each of the 16 carbon sources in single-resource environments (Fig. 1b). In our model, for simplicity, each species consumed one resource to grow, to approximate particular resource preferences in different species. It then leaked metabolic byproduct(s) into the environment, each of which was one step downstream from the consumed metabolite according to the cross-feeding network (see Methods). Other species could then consume these leaked metabolites, in turn releasing new by-products into the environment (Fig. 3b). Importantly, leaked byproducts always comprised a fixed fraction of the consumed resource, resulting in a progressive decrease in the concentration of metabolic byproducts available to microbes downstream^34,35^ (Fig. 3b). Finally, to account for metabolic overflow^51–53^, we added a small quantity of TCA intermediates and acetate to all simulated media. Overall, this model incorporated ecological dynamics and cross-feeding in a realistic fashion while retaining simplicity.

**Figure 3.**
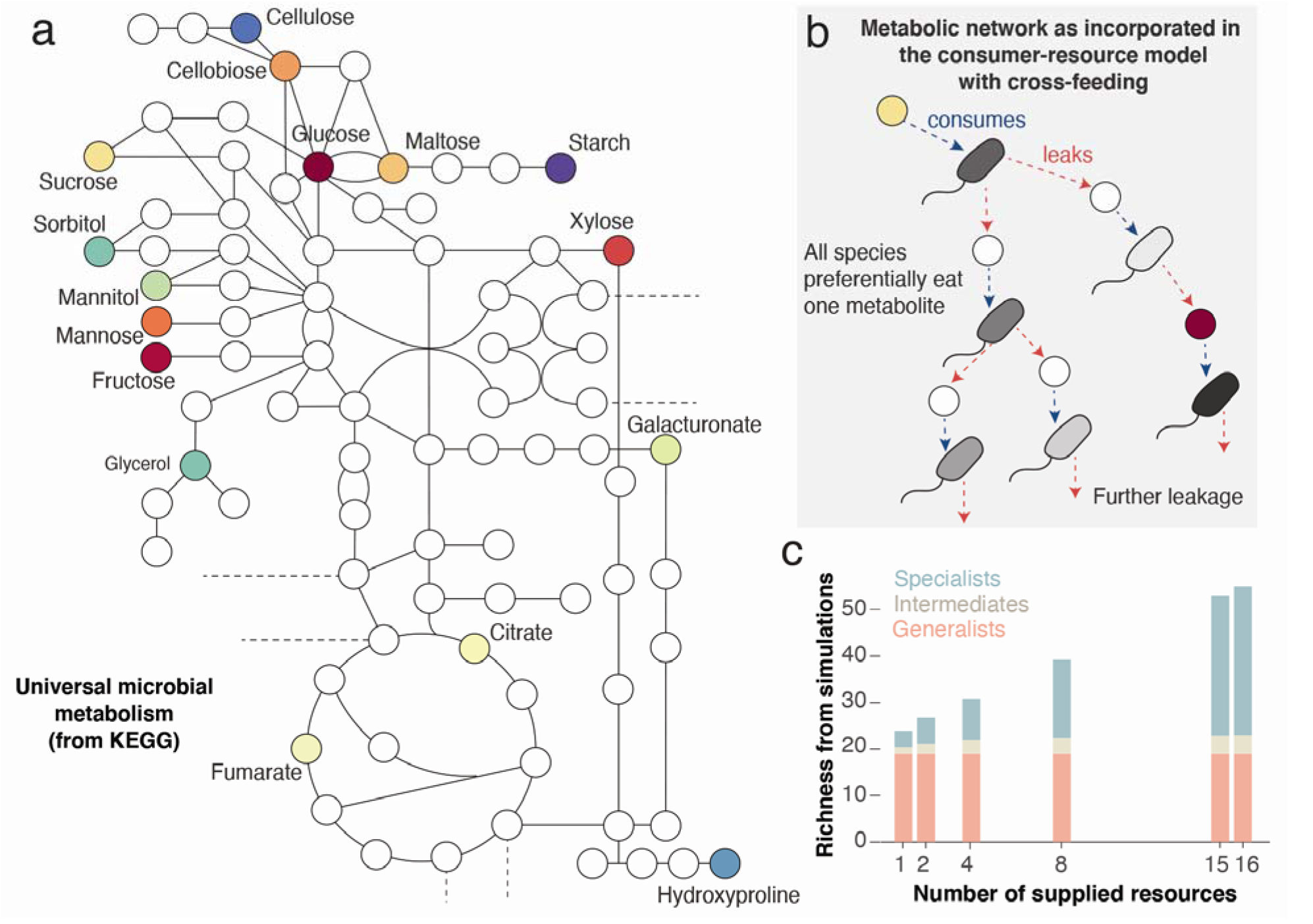
A resource explicit model incorporating a realistic metabolic network recapitulates our experimental results. **a**. A simplified version of the metabolic map derived from KEGG and MetaCyc databases is shown, where the carbon sources used in the experiments are highlighted. The metabolic map is used to build the cross-feeding matrix used in the model. **b**. Schematic showing the flow of metabolic byproducts in our model. Colored circles indicate supplied resources; white circles indicate metabolic byproducts, i.e., metabolites that are downstream from the resource in the metabolic network; colored microbes indicate different microbial taxa; and arrows indicate leakage of metabolic byproducts, which serve as resources for other taxa. **c**. Richness obtained from simulations with all the possible combinations of 16 resources (14,843 conditions in total) is plotted as stacked bars indicating the average number of species for each category: generalists (pink), specialists (teal) and intermediates (beige). Species in the model are classified based the number of “media” they survived in, analogously to the distinction applied in the experiment (Fig. 2). Error bars are omitted for clarity. The total richness increases linearly with the number of resources (intercept = 22.7, slope = 2).

Simulations of this model reproduced our two most-prominent experimental observations. That is, we could observe the stable coexistence of many species in single resources (between 19 and 28 species, Fig. S16) yet a modest linear increase in richness with the number of resources (slope ∼2, Fig. 3c, Fig. S16). Since the estimated metabolic byproducts came from mapped metabolic reactions, we already knew that their number increased non-linearly with supplied resources (Fig. S9). So, how could we get a linear resource-diversity relationship? In our simulations, the concentration of byproducts varied with two factors: their position in the metabolic network and the initial concentration of the supplied resources. To mimic our experiment, we maintained a constant total resource concentration. This resulted in the concentration of each supplied resource decreasing with the total number of supplied resources in the medium (as 1/*R, R* being the number of supplied resources). As we provided more resources, a progressively larger fraction of byproducts had their steady-state concentrations decrease non-linearly. Byproducts at very low concentrations could no longer support microbial species, since their growth rates fell below the dilution rate. Hence, even though the number of metabolites grew non-linearly with the supplied resources, the metabolites that could support the growth of new species grew much slower, resulting in a linear increase in diversity in our simulations.

By classifying the species in our simulations based on their resource occupancy (as in the experiment), our model also predicted a constant number of generalists and an increasing number of intermediates and specialists with additional resources (Fig. 3c). This is consistent with our experimental observations (Fig. 2g), and further corroborates the idea that the generalists observed in our communities were better at taking advantage of core metabolites, while the specialists that survived were those that were better at competing for rarer metabolites.

Importantly, by implementing our model with a realistic network, we were able to simulate all the possible combinations of our sixteen resources, even those that we did not experimentally grow. These simulations showed the same relationship between richness and resource availability (Fig. 3c), suggesting that the outcome of our experiment would not have changed if we had included more/different resource combinations. We concluded that the realistic cross-feeding network seeded by the pool of carbon sources in our experiment could explain the observed relation between microbial community diversity and resource number.

## Discussion

Understanding the relationship between available nutrients and community diversity is central to both theoretical and experimental ecology. Here, using a high-throughput culture enrichment approach amenable to mathematical modeling, we provide experimental and theoretical evidence of how the identity and the number of available resources modulate microbial community diversity via a network of metabolic cross-feeding interactions. We showed that the richness of communities grown on single sources of carbon can be predicted from the number of cross-fed byproducts generated using intracellular metabolic reactions fueled by those resources. In addition, using this realistic metabolic network as the cross-feeding network in a resource-explicit model was sufficient to reproduce the observed linear, modest increase of richness with the number of available resources.

Our results add to the wealth of studies stressing the importance of metabolic cross-feeding as a pivotal driver of species coexistence^54^ and its link to the identity of available resources^18,27^. We have observed multispecies communities on all compounds provided as single sources of carbon, including mono-, di- and polysaccharides, sugars alcohols and organic acids. Importantly, we were able to link the variability in community richness to the identity of the supplied resource by mapping the metabolic pathways triggered by each resource and estimating the number of byproducts potentially produced and leaked into the growth media. We are aware that these predictions do not provide any information on the bio-availability of the metabolic byproducts and might be biased towards well-characterized bacterial species. Nevertheless, they provide a simple and tractable way to estimate byproducts using only the structure of an overall metabolic network. Incidentally, community-scale flux balance simulations on the single-resource communities in our experiment also predicted a correlation between the number of expected byproducts and community richness (Fig. S17 and Methods). Further advances in linking the available byproducts with richness could be provided by targeted metabolomics, a technique which can assess the relative concentration of the metabolites in a medium, as in ref.^51^.

Together with the remarkable richness in single resources, the other striking characteristic of our results was the modest linear increase in community diversity with the number of additional nutrients. Both these features appeared to stem from the structure of the metabolic network seeded by the pool of resources included in the experiment. Indeed, implementing this realistic network in a resource-explicit model was sufficient to recapitulate both features. Another important ingredient of the model was the concentration of metabolites, which in turn depended on their position in the metabolic network and the concentration of the supplied resource they were generated from. Despite its simplifying assumptions (e.g., we used species-independent growth and leakage rates), our model captured the combined effect of dilution and resource concentration that might have determined the diversity of our experimental communities. At the same time, while simulations of our model recapitulate the observed relationship, its theoretical bases still remain to be fully understood, including an extensive exploration of how the structure of the metabolic network affects resource-diversity relationships. Other approaches to complement such theoretical efforts might include experimentally testing the effect of resource concentration and quantitatively modelling intracellular metabolism, thus also accounting for metabolic fluxes and redox balances^55–59^.

The position from which a resource enters the central metabolism affects not only its availability but also the direction in which metabolic reactions run (i.e., glycolytic or gluconeogenic). We showed that whether a resource is glycolytic or gluconeogenic was an important predictor of the diversity and structure of microbial communities, as it dictated the ratio between habitat generalists and specialists. These results suggest, and a two-parameter regression supports (Fig. S18), that adding gluconeogenic resources (e.g., organic acids) while keeping the total concentration of carbon constant may not increase the community diversity. Overall, our results add to the studies stressing that the position from which a resource enters the central metabolism eventually determines its use, including diauxic shifts vs. co-utilization^60^ and tradeoffs between growth and lag in changing environments^51^.

A further indication of the role played by the metabolic network sustained by the supplied resources came from the striking parallelism that we observed between the structure of experimental communities and the architecture of the network itself. Just like metabolism consists of shared (e.g., the TCA cycle) and unique reaction modules (i.e., specific to the degradation of a particular resource), all experimental communities harbored a core group of metabolically flexible, faster-growing habitat generalists and variable numbers of taxa associated with a particular nutrient (habitat specialists). This suggested that the habitat generalists present in our stabilized microbial communities were “specialists for common nutrients”, i.e., they preferentially consumed substrates that are commonly produced during bacterial growth. In this sense, generalists growing on downstream metabolites (e.g., TCA intermediates) depended on specialists for the production of their favorite substrates. Consistent with this, community flux balance simulations where we paired a generalist and specialist ASV showed that it was the specialists that are likely to leak metabolic byproducts used by generalists, and not *vice versa* (Fig. S19 and Methods).

Finally, in our experiments, habitat specialists outnumbered generalists on the whole, a pattern that is commonly observed when natural communities from different locations are compared^38^. Surveys of microbiomes across different ecosystems have also highlighted a remarkable level of determinism in the association between microbiome composition at coarse taxonomic resolutions (e.g., at the family-level) and availability of nutrients. This feature is recapitulated by other studies^18,61^. Here we showed both the persistence of strong taxa-resource associations at the ASV (Fig. 2f) and the family level, with the relative abundance of several families, comprising prevalently either generalist or specialist taxa, changing as a function of the relative concentration of specific resources (see Methods for how we established which resources influenced the most each family). Specifically, we observed that the relative abundance of several specialist families decreased drastically or went to zero when the relative concentration of the “favorite resource” dropped by half (e.g., Cellvibrionaceae, Fig. S20), while the relative abundance of generalist taxa increased non-linearly with the relative concentration of few resources, e.g., Pseudomonadaceae with hydroxyproline and fumarate (Fig. S20). The fact that empirically observed features of natural microbial communities emerge in controlled experiments suggests that they might reflect the effects of deterministic processes linked to nutrient availability rather than be generic emergent properties of complex multi-agent systems.

## Methods

### Growth media preparation

All the chemicals were purchased from Sigma-Aldrich unless otherwise stated.

All bacterial cultures were grown in M9 media (prepared from 5X M9 salts, 1X Trace Metal Mixture (Teknova) and 1M stock solutions of MgSO_4_ and CaCl_2_) supplemented with 0.1 % w/v of one of 75 carbon source combinations. These combinations include: 16 compounds commonly available in soil that were provided as single carbon sources (D-(+)-glucose, D-(–)-fructose, D-(+)-xylose, D-(+)-mannose, D-(+)-cellobiose, D-(+)-maltose monohydrate, sucrose, citric acid, fumaric acid, D-(+)-galacturonic acid monohydrate, D-mannitol, D-sorbitol, glycerol, trans-4-Hydroxy-D-proline, methyl cellulose, starch); 24 random combinations of two of these resources; 12 random combinations of four resources; 6 random combinations of eight resources; the 16 combinations containing 15 resources; and all the 16 resources together (see Table S1 for the complete list and Fig. S2). The total concentration of carbon was kept the same and resources were in all instances supplied in equal amounts, that was 100%, 50%, 25%, 12.5%, 6.7% and 6.25% each for single-, two-, four-, eight-, 15- and 16-resource combinations. All solutions were filter-sterilized with a 0.22 *μ*m filter and kept at 4°C for the duration of the experiment.

### Collection of microbial communities from the environment

The soil from which the initial inoculum comes from was sampled from a lawn in Cambridge, Massachusetts, at a depth of ∼15 cm using a sterile corer and tweezers. Once in the lab, a total of 1.5 g of the collected soil was diluted in 20 mL phosphate buffered saline (PBS; Corning), then vortex at intermediate speed for 30 s and incubated on a platform shaker (Innova 2000; Eppendorf) at 250 r.p.m. at room temperature. After 1 hour, the sample was allowed to settle for ∼5min and the supernatant was filtered with a 100 *μ*m cell strainer (Thermo Fisher Scientific) and then directly used for inoculation. Both the original soil sample and the remaining supernatant were stored at −80 °C for subsequent DNA extraction.

### Experimental microcosms

Aliquots (7*μ*L) of the supernatant containing the soil microbial suspension were inoculated into 203 *μ*L of growth media in 96-deepwell plates (Deepwell plate 96/500 *μ*L; Eppendorf), for a total of 231 microcosms (3 replicates for each different resource combinations, except 16-resource combinations that were replicated 9 times). Deepwell plates were covered with AeraSeal adhesive sealing films (Excel Scientific). Bacterial cultures were grown at 30°C under constant shaking at 1,350 r.p.m. (on Titramax shakers; Heidolph). To avoid evaporation, they were incubated inside custom-built acrylic boxes.

Every 24 h, the cultures were thoroughly mixed by pipetting up and down 3 times using the VIAFLO 96-well pipettor (Viaflo 96, Integra Biosciences; settings: pipette/mix program aspirating 7 *μ*L, mixing volume 10 *μ*L, speed 6) and then diluted 1/30x into fresh media. We applied a total of seven daily dilution cycles. At the end of every cultivation day we measured the optical density (OD_600_) using a Varioskan Flash (Thermo Fisher Scientific) plate reader. The remaining bacterial culture was frozen at −80 °C for subsequent DNA extraction.

### DNA extraction, 16S rRNA sequencing and analysis pipeline

DNA extraction was performed with the QIAGEN DNeasy PowerSoil HTP 96 Kit following the provided protocol. The obtained DNA was used for 16S amplicon sequencing of the V4 region. Library preparation and sequencing, which was done on an Illumina MiSeq platform, were performed by the MIT BioMicroCenter (Cambridge, Massachusetts).

We used the R package DADA2 to obtain the amplicon sequence variants (ASVs)^62^ following the workflow described in Callahan et al.^63^. Taxonomic identities were assigned to ASVs using the SILVA version 132 database^64^. The phylogenetic tree (Fig. S4) was reconstructed using Randomized Axelerated Maximum Likelihood (RAxML) using default parameters^65^.

### Data analysis

Analysis, unless otherwise stated were conducted in R, version 3.6.1^66^.

Sequencing data was handled using the R package phyloseq^67^. We obtained an average of 20,613 reads per sample. Sequencing depth did not affect our estimation of community diversity indexes (Fig. S4). Richness was calculated as the number of ASVs with abundance larger than 0 found in each sample. Community diversity was also measured by Shannon Diversity index and Shannon Entropy index following^68,69^ (Fig. S10). The significance of differences in richness due to single supplied resources was tested through ANOVA^70^ using the package GAD.

### Richness predictions

Predictions of how richness would grow with the number of supplied carbon sources were computed using all the three replicated communities grown on a single resource and all the possible combinations of single resources (120 combinations of two resources, 1,820 combinations of four, 12,870 combinations of eight, 16 combinations of 15 and one combination of 16 resources) (Fig. 1D). As an example of the prediction based on the maximum of constituent singles, the richness of the community grown in a medium containing glucose + hydroxyproline was obtained by calculating the maximum richness over each couple of replicates (one containing only glucose and the other containing only hydroxyproline) and subsequently averaging across all the predicted maxima (in total 9 predicted values). The same procedure was used for the average of constituent singles. Analogously, for the predictions based on the union of constituent singles, the richness in glucose + hydroxyproline was predicted by calculating the number of unique ASVs found in each couple of replicates of constituent singles (i.e., the total number of ASVs minus the number of overlapping ASVs) and then averaging across all obtained unions (9 values).

### Rank abundance distributions

First, we computed abundance distributions (RADs) for each sample, i.e., each replicate community grown on a unique combination of carbon sources, by sorting ASVs based on their relative abundance. Then, we plotted the RADs in a log-linear fashion and fitted a regression line in order to compare their slopes (Fig. S11A). The absolute value of the slope of the fitted regression line informs on the abundance distribution of the ASVs in a community. More even communities usually display smaller slopes (Fig. S11B). Since each community exhibited a different richness, we normalized the RADS for richness (Fig. S11C) To do this, we used the *RADnormalization_matrix* function in the RADanalysis package: from each RAD with an observed richness, this function generates a “normalized RAD” with a richness corresponding to the minimum richness observed in the experiment (7 ASVs) by randomly resampling the original RADs for 10 times^71^. In this way, samples with different richness can be compared and changes in evenness properly assessed.

### Definition of generalists and specialists based on single resource occupancy

ASVs found in single resources were classified in three categories based on how many media containing a single resource they were found in, i.e. they exhibited abundance larger than 0 ^38,43^. We considered specialists the ASVs that were observed in less than 25% of single-resource media, i.e., in one, two or three resources. Generalists were those ASVs found in more than the 75% of media, i.e., in 13 or more resources. We defined intermediates the ASVs found between four and twelve resources. These thresholds were chosen arbitrarily, but the resulted in about ∼ 4% generalists and 80% specialists, consistently with proportions of generalists and specialists observed in natural communities ^38,39,43^. We chose this simple way of assigning ASVs to generalist, intermediate and specialist categories over other methods, e.g. as in ^24^ in order to leave aside their relative abundance, which was analyzed separately.

### Prediction of possible metabolic byproducts in resource environments

We predicted the possible number of metabolic byproducts that could be produced using the resources present in each medium using a curated metabolic network. The metabolic network contained a large set of metabolic reactions encompassing carbohydrate, sugar and amino acid metabolism extracted from the KEGG database^31^. We manually curated this large set of reactions using the MetaCyc database^32^ in order to limit it to reactions possible by most microbial taxa common to the soil, such as *Pseudomonas*. We used this network to estimate all the metabolic compounds that could be produced as byproducts, starting from the carbon sources available in each medium. We assumed that a small set of “currency” molecules, such as water, carbon dioxide and ATP, were always available as reactants when required (full list of currency molecules: phosphate, oxygen, carbon dioxide, water, H^+^, ATP, NAD(P)H, Acetyl-CoA, CoA).

To estimate the possible byproducts in each medium, we employed the well-known scope expansion algorithm^72–76^. Each reaction in our curated metabolic network consisted of a set of reactants and resulting products. For each medium, we first asked which reactions could be performed using only the carbon sources available in the medium (i.e., the current “scope” of the medium). We assumed that the products of these reactions could be produced and added them to the set of reactants – the new scope – for the next step. In the next step, we again asked which reactions could be performed using the new scope. We added their products to the scope for the next step. We continued this process, step by step, until we could add no new products to the scope. The resulting final scope of metabolites, minus the currency molecules provided in the medium, was our estimated set of possible metabolic byproducts producible in that medium.

Adding some amino acids as currency molecules, mimicking our experimental protocol, yielded a larger set (∼3x) of possible metabolic byproducts for each medium, including many amino acids and anabolic products. This expanded set of metabolites for each medium was also correlated with the observed average species richness in that medium (data not shown).

We also tried an alternative approach to estimate the number of metabolic byproducts in single resource environments, using community-scale flux balance simulations. For each ASV observed in a single resource environment, we first obtained the phylogenetically closest whole genome sequence in NCBI’s RefSeq database. For this, we mapped the 16S sequence of each ASV to complete genomes in the RefSeq database using BLAST^77^. For each ASV, we chose the genome that had the highest identity; when multiple genomes matched this criterion, we chose the longest genome, following similar work^78^. We then obtained all the mapped genome sequences and constructed metabolic models for each of them using CarveMe^55^; we gap-filled all models to grow on M9 minimal medium supplemented with metal ions, such as iron and copper, which are present in trace amounts in experimental bacterial growth media.

To estimate the number of metabolic byproducts in each single resource environment, we performed community-scale metabolic simulations using the package MICOM^57^. For each community, we input all the metabolic models for all ASVs detected in that community, and simulated their growth in the corresponding media. We then counted all metabolites which were predicted to be exported by each community as the estimated number of byproducts for that community. For each medium, we averaged the number of byproducts across all three replicate communities; we used this as our estimated number of byproducts for that medium.

### Characterization of the structure of the metabolite pool

Following the same logic that we used for ASVs, metabolites estimated to be produced through metabolic reaction starting from single resources were classified in three categories based on the number of resources that could be produced from. We considered *rare* metabolites those observed in less than 25% of single-resource media, i.e., in one, two or three resources. In contrast, *common* metabolites were those found in more than the 75% of media, i.e., in 13 or more resources. Finally, *intermediate* metabolites were those present in between four and twelve resources. The chosen thresholds separate the metabolites of the central metabolic pathway (common metabolites) from the peripheral metabolites belonging to branches descending into the central pathway (rare and intermediate metabolites).

### Inference of rRNA operon copy number for generalist and specialist taxa

To test for signatures of different life-history strategies of the generalist and specialist taxa in our study, we estimated their 16S rRNA operon copy numbers. We estimated rRNA copy numbers at the level of both genus and family, separately for generalist and specialist taxa. For each genus identified, we queried rrnDB^79^—a database of rRNA operon copy number statistics—for the median copy number corresponding to the genus. We used this as an estimate for the rRNA operon copy number of that genus.

### Inference of number of metabolic genes for generalist and specialist taxa

To test for metabolic differences between the generalist and specialist taxa in our study, we estimated the number of metabolic genes in their genomes. Since we did not have either isolates or assembled genomes corresponding to the observed taxa, we relied on a popular indirect method of estimating gene content. Namely, for each ASV, we used the reference genome which was phylogenetically closest to that ASV as a proxy for its genome. For this, we used PICRUSt2^80^ using default parameters; as an input to the tool, we provided the 16S rRNA sequences of all 226 generalist and specialist taxa as well as their abundances in each sample. After running PICRUSt2, we obtained a table of the predicted gene content for each ASV (i.e., presence/absence of a specific KO number in the KEGG database). We extracted all metabolic genes from this table by only choosing those KO numbers which had at least one known metabolic reaction corresponding to them. Doing so resulted in an estimated set of metabolic genes for each ASV; we used this as an indirect estimate of the metabolic capabilities of each ASV.

### Calculation of the resource-specificity score

We used a resource-specificity score to test if the ASV-resource associations that we observed in single resources were maintained when the single resource(s) in which the ASV was found was combined with others. For each ASV present in a single resource (target resource), the resource specificity score is calculated as the difference between the number of multi-resource media containing the target resource in which the ASV is found and the number of media not containing the target resource in which the ASV is found divided by the total number of media in which the ASV is found (Fig. 2E). This is reminiscent of a preference index, which is a standard measure in the behavioral sciences. Single resources are excluded from the count. The resource-specificity score ranges from 1, indicating that the ASV is present only when the target resource is provided, to −1, implying that the ASV is always absent when that resource is supplied with other resources. A score of 0 is indicative of an ASV showing no specificity for that particular resource (Fig. 2E). We calculated a score for each ASV-resource pair, such that each ASV had as many scores as the number of single resources is found in. Then, we computed the average of the scores obtained for each single resource, separating between scores belonging to generalist and specialist ASVs (Fig. 2E).

### Inference of metabolic interactions between generalist and specialist taxa

To estimate whether metabolic interactions between the generalist and specialist taxa in our communities were likely to be unidirectional or bidirectional, we used SMETANA v.1.0^56^, using default settings. For each generalist-specialist pair that we experimentally detected in single resource environments, we used SMETANA on a model community comprising both ASVs (a generalist and a specialist) using the settings --flavor bigg --exclude inorganic.txt -d. We explicitly disallowed inorganic molecules such as phosphates, carbon dioxide and metal ions from being exchanged by using the –exclude option in SMETANA. To consider interaction directionality, we looked at the donor and receiver of each exchanged metabolite. When there was only one donor for every exchanged metabolite, we inferred the interaction as unidirectional, with the direction going from the donor to the receiver of the metabolites.

### Detection of family-resource associations using an ensemble tree regression model

We calculated the relative abundance of the most prevalent families (37) in the 75 replicated bacterial communities and ran an ensemble tree regression model to detect significant patterns of variations in family abundance due to changes in the relative concentration of resources.

We chose to coarse-grain the abundance data at the family level because, while several ASVs were lost and others were gained going from one to 16 resources in the growth media, the families found across all combinations of resources were mostly the same. In addition, we distinguished between generalist families, i.e., those that contained at least one generalist ASV, and specialist families, i.e., containing only specialist ASV. Consistent with ASV-level definition, generalist families displayed higher mean rRNA operon copy number compared to specialist families.

We employed XGBoost, a gradient boosting framework based on decision trees^81^. Specifically, we implemented a regression model for each family in which the input was the relative resource concentration and the output was the log-transformed relative family abundance. We trained the model on two replicates by performing leave-one-out crossvalidation of the XGBoost parameters “max_depth”, “n_estimators” and “learning_rate”^82^ and tested on the third one with average mean-squared error across families of 6.05. We applied the Shapley Additive exPlanations (SHAP)^83^ to identify the resources that were more important in driving changes in the abundance of each family. This analysis has been done using Python version 3.8.

Results of this analysis revealed that variations in the abundance all of the 37 families were driven by one or multiple resources based on their dominant life strategy. To simplify the visualization of the results we plotted the relative abundance of some representative families as a function of the concentration of the resources identified by the analysis (Fig. S20). Families mostly composed of specialist taxa, e.g., Cellvibrionaceae and Bacillaceae, showed abrupt changes in their abundance with the concentration of the “favorite” resource (Fig. S20). By contrast, more generalist families, e.g., Pseudomonadaceae and Enterobacteriaceae, exhibited smooth trends in their abundance with the concentration of multiple resources.

### Resource-consumer model with cross-feeding and simulations

The parallelism between species and metabolite distribution (see Fig. 2) that we observed in our experiment highlighted that the cross-feeding network is key to understand microbial communities under each combination of supplied carbon sources. To test this idea, we used a model encompassing the metabolic network that we obtained from the analysis of KEGG and MetaCyc databases. This was achieved by a consumer-resource model with cross-feeding^18,21,35^. In our consumer-resource model with the realistic metabolic network, we made the following simplifying assumptions.

First, we assumed that every species consumes only one preferred metabolite. Upon this assumption, competitive exclusion guarantees that only the best grower in each resource survives; thus, we implemented only one species for each resource in our simulation as a post-selection pool. This assumption reflected the resource-species association we observed (Fig. 2), which suggested that the taxa identified as generalists may specialize on core metabolites that are found everywhere. Also, while many species can consume multiple resources, they may still grow much faster on the most preferred one. Metabolic strategies such as *diauxie* also highlight that growth on the most preferred resource can be a dominant factor for community assembly^51^.

Second, growth rates, biomass yield, and leakage rates (these quantities are described below) are universal, independent of species identity. This assumption led to the simplest implementation of our metabolite network.

Third, we assumed that each species leaked out all the immediate metabolites of the metabolite it consumes. The list of immediate metabolites that are produced from each metabolite was obtained from scope expansion analysis. This information is encoded by a cross-feeding matrix *CF*_*ij*_, which is nonzero when *i*^*th*^ metabolite immediately leaks *j*^*th*^ metabolite and 0 otherwise. For simplicity, the nonzero values of *CF*_*ij*_ are set to be 1/(number of metabolites produced by *i*^*th*^ metabolite).

The scope expansion analyses based on the metabolic reactions mapped in KEGG and MetaCyc databases identified 96 metabolites that could be produced starting from the supplied carbon sources. Thus, CF is a 96×96 matrix. The original scope expansion analysis included reactions where multiple reactants were required to generate products. Since it is impossible to fully capture such interdependences with a matrix, we assumed that reactions were activated as long as one or more reactants were present. Also, to mimic the highly connected and cyclic structure of TCA-cycle, we set each TCA intermediate to generate all other TCA intermediates.

Under these assumptions, we simulated the dynamics of the following model:

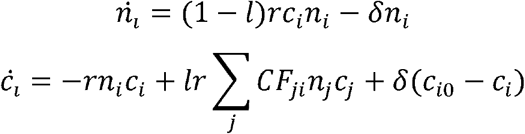

where *N*_*i*_ is the population of *i*^*th*^ species, and *c*_*i*_ is the concentration of *i*^*th*^ metabolite. is the leakage rate, *l* is the per-capita, *r*_*i*_ per-resource growth rate of *i*^*th*^ species, delta is the dilution rate of the chemostat-like environment. *CF*_*ij*_ tells whether *i*^*th*^ metabolite is leaked from *j*^*th*^ metabolite based on the scope expansion analysis. *c*_*io*_ s the supply resource concentration corresponding to each combination of supplied carbon sources, controlled by overall scale *c*_*o*_. For example, when glucose is supplied, *c*_*io*_ *= c*_*o*_ for *i*=glucose and 0 otherwise. And when a combination of glucose and hydroxyproline is supplied, 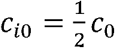 for *i*=glucose, hydroxyproline and 0 otherwise. To simulate the effects of metabolic overflow^51–53^, we supplied a small quantity (c = 0.2) of TCA intermediates and acetate to all media.

The first equation models the dynamics of population. The first term tells that the growth rate of each species is proportional to the concentration of the preferred resource. We also assumed that species can only convert a fraction 1 − *l* of the preferred resource into biomass, since *l* is leaked in the environment as by-product(s). The second term represents dilution as the main driver of mortality in this chemostat-like system.

The second equation models the dynamics of resources (both supplied and cross-fed). The first term represents the consumption of the resource by the specialized species. The second term represents the leakage from upstream resources that cross-feed *i*^*th*^ the resource. The third term represents the dilution and external supply of resource in the chemostat system.

We simulated the model dynamics under all possible combinations of 1, 2, 4, 8, 15, and 16 number of supplied resources (14843 combinations total). We chose the parameters *δ* 0.1, which is comparable to the dilution we imposed in the experiment, *r* = 1, *c*_*o*_ = 100, and *l* = 0.1. In Fig. 3c we show the results of all combinations, while in Fig. S16 we plotted only the combinations included in the experiment. The simulations were run for 1*e*^3^ unit time starting from initial population set as *e*^−7^, and communities reached equilibrium at the end of the simulations. The population cutoff for survival was set as *e*^−7^. Simulations were run in Python version 3.7.4.

## Data availability

Data files and analysis/simulation codes will be available via GitHub upon publication. 16S Amplicon sequencing data and metadata files have been deposited in the NCBI SRA database under NCBI BioProject ID PRJNA715195.

## Acknowledgements

The authors would like to thank Jacopo Grilli, Marco Costantino-Lagomarsino, Mattia Corigliano and Matthieu Barbier for feedback on models, the members of Gore Lab for comments on the manuscript, and Bartolomeo Stellato for the help with the ensemble tree regression model. This work was supported by NIH and the Schmidt Foundation. A.G. is supported by the Gordon and Betty Moore Foundation as a Physics of Living Systems Fellow through grant number GBMF4513.

## Author contributions

MDB and JG conceived the study. MDB performed the experiments and the sequencing analysis. HL performed theoretical modeling. AG performed metabolic and genomic analyses. All authors analyzed the data and wrote the manuscript.

## Materials & Correspondence

Data files and analysis/simulation codes will be available via GitHub upon publication and can be requested to. 16S Amplicon sequencing data and metadata files have been deposited in the NCBI SRA database under NCBI BioProject ID PRJNA715195.

