## Supplementary Figures and Tables for "Resource-diversity relationships in bacterial communities reflect the network structure of microbial metabolism"

**This file includes:**

Figs. S1 to S20

Tables S1

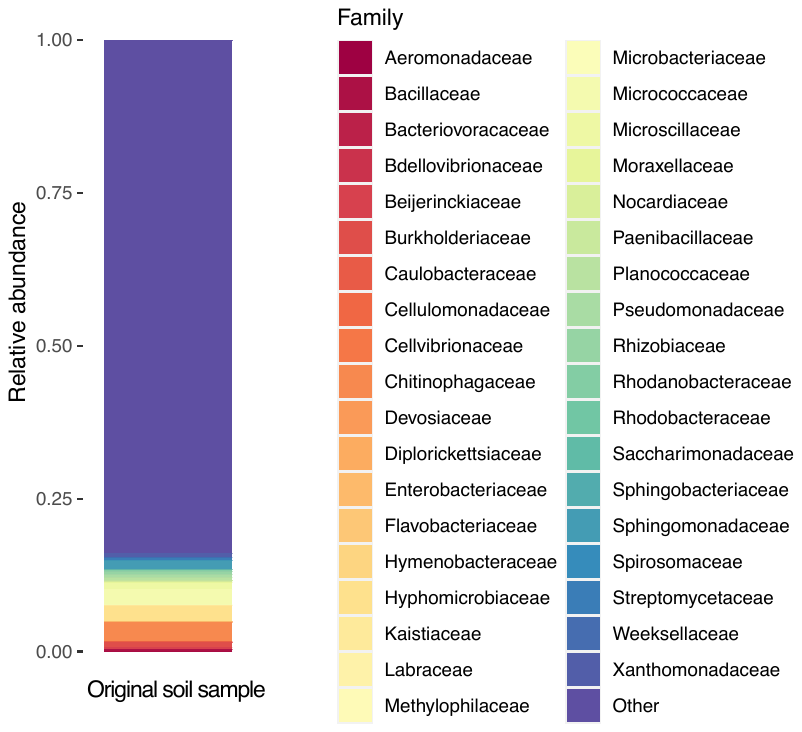

**Fig. S1. The soil sample used to inoculate the experimental microcosms is highly diverse and taxonomically rich.** A total of 750 ASVs were detected in the soil sample using 16S rRNA amplicon sequencing at single nucleotide resolution Here the relative abundance of the 39 most prevalent families in the experimental microcosms after 7 days is shown. These families constitute less than the 20% of the relative abundance of soil taxa.

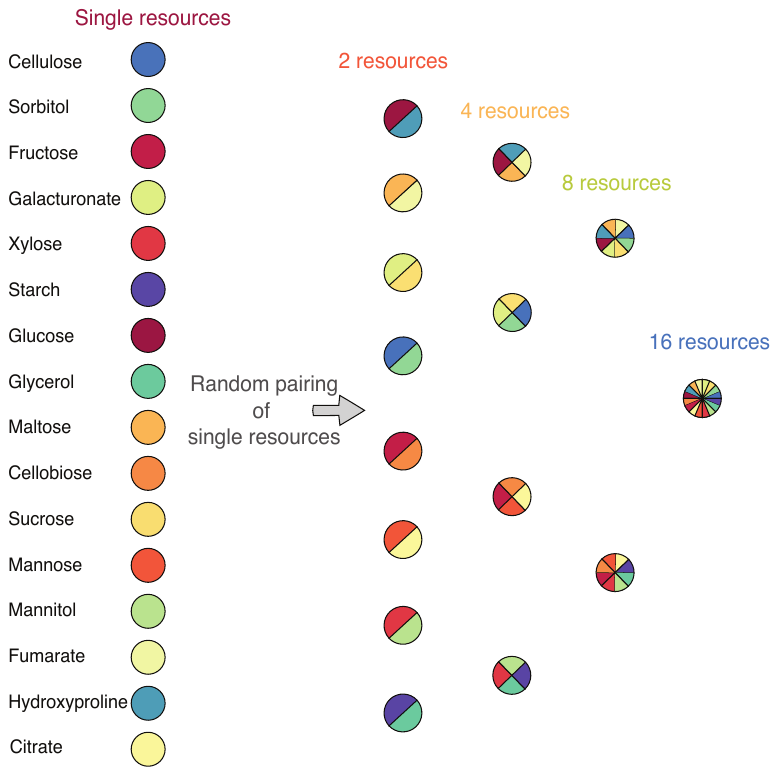

**Fig. S2. Example of random grouping of single resources in 2, 4, 8 ,16-resource combinations.** A total of three random groupings were included in the experiment. The total carbon concentration is kept the same across different combinations of resources (0.1 % w/v). Also, all 15-resource combinations were included.

**Fig. S3. The majority of communities reached equilibrium before the end of the experiment.** Each panel shows the temporal trajectories of the composition of one community at the family level. The most prevalent 37 families are included. The first 16 plots depict communities grown on a single carbon source. The last four plots depict replicated communities grown on a media containing all the 16 carbon sources.

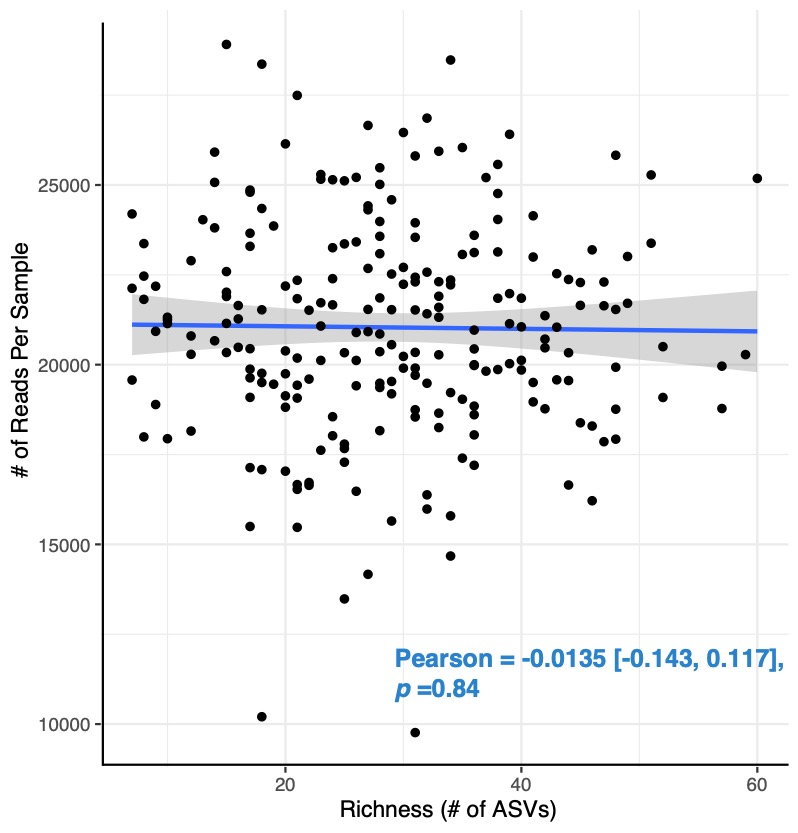

**Fig. S4. Sequencing depth did not affect richness estimates.** There is no correlation between the richness of a sample and the number of reads obtained for that sample (number of samples 227). The estimated Pearson correlation coefficient is not different from zero.

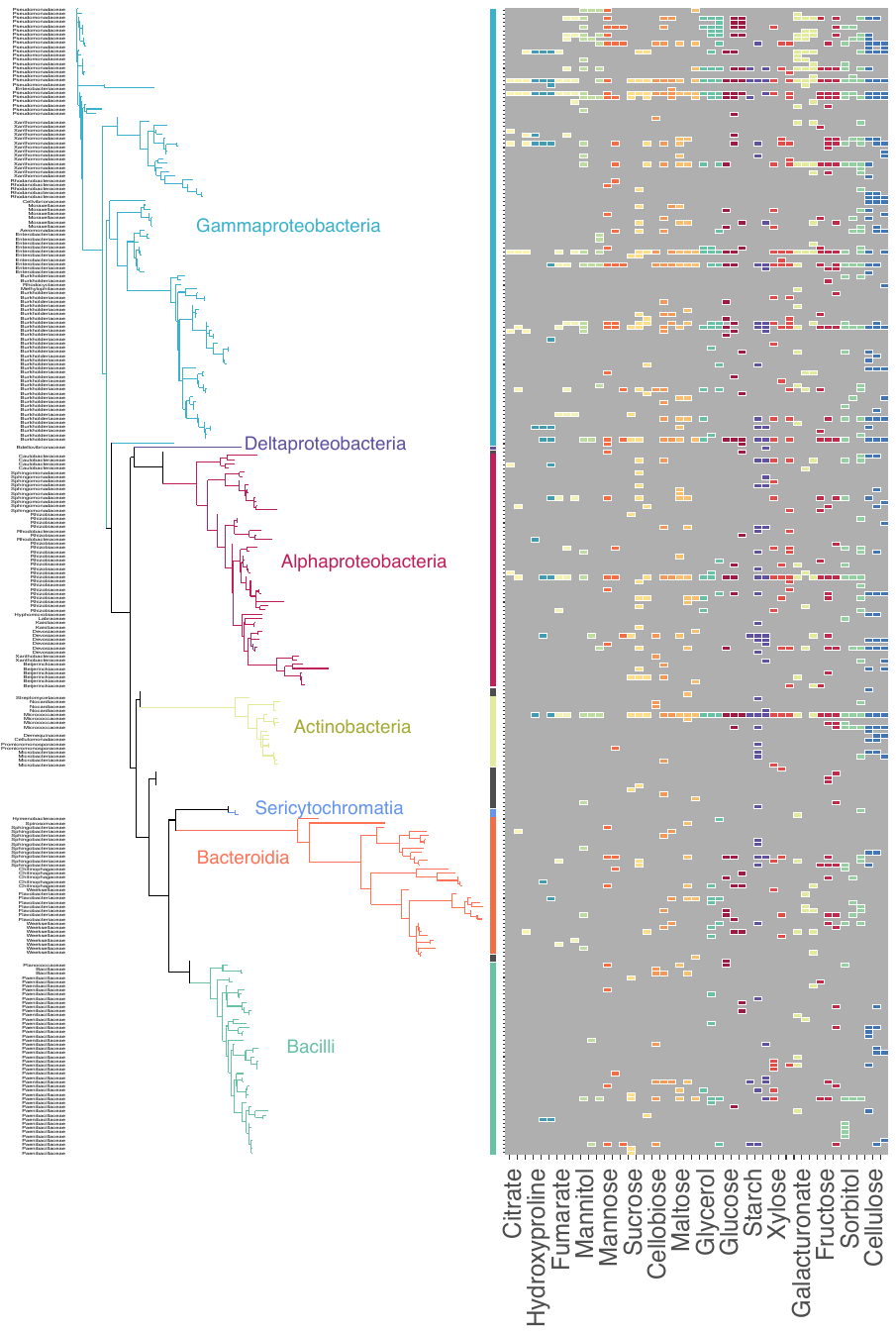

**Fig. S5. Single carbon sources support microbial assemblages spanning a wide phylogenetic diversity.** The pool of ASVs found across all media supplied with single resources is phylogenetically diverse, encompassing 7 classes, indicated by colored lines on the right side of the plot (black lines indicate ASVs that could not be identified at any taxonomical level except the Domain, Bacteria). Families are indicated on the left side of the phylogenetic tree. Colored tiles indicate the media in which ASV is found (for each carbon source, there are three replicated microcosms for a total of 48 communities.) Carbon sources are ordinated in an increasing order based on the average richness they support.

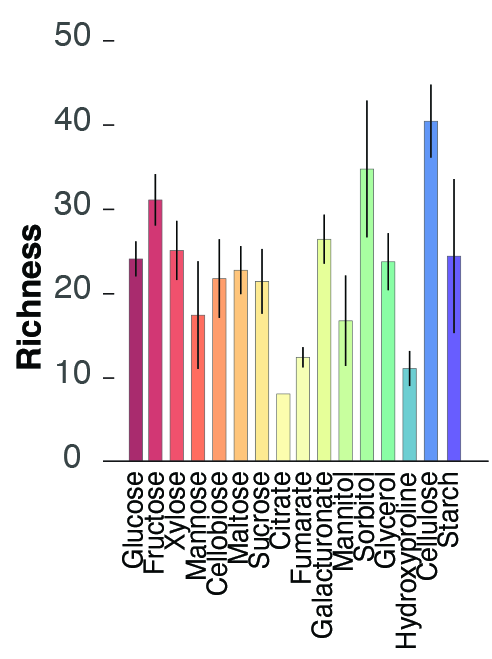

**Fig. S6. All individually-supplied carbon sources supported multispecies communities, but richness varied with the identity of the resource.** Bars indicate, for each carbon source, the number of ASVs (mean $\pm$ SEM, N = 3).

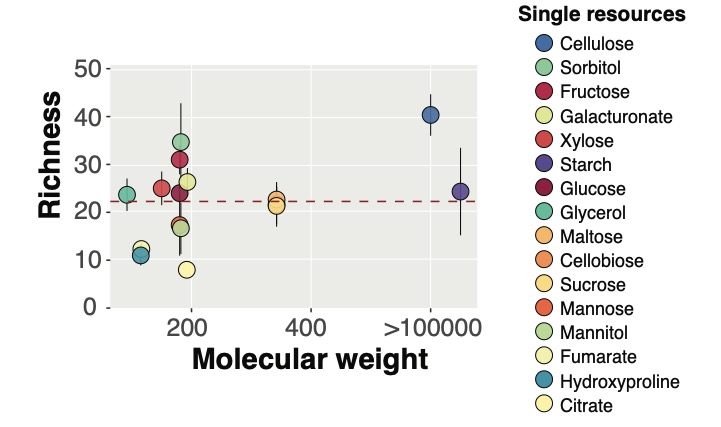

**Fig. S7. Richness of microbial communities in single resources is not explained by the molecular weight of the supplied resource.** MW of cellulose and starch is reported as > 100,000 for graphical reasons. The red dashed line represents the estimate linear regression including the actual MV of cellulose and starch. The color of the dots indicates the supplied resource.

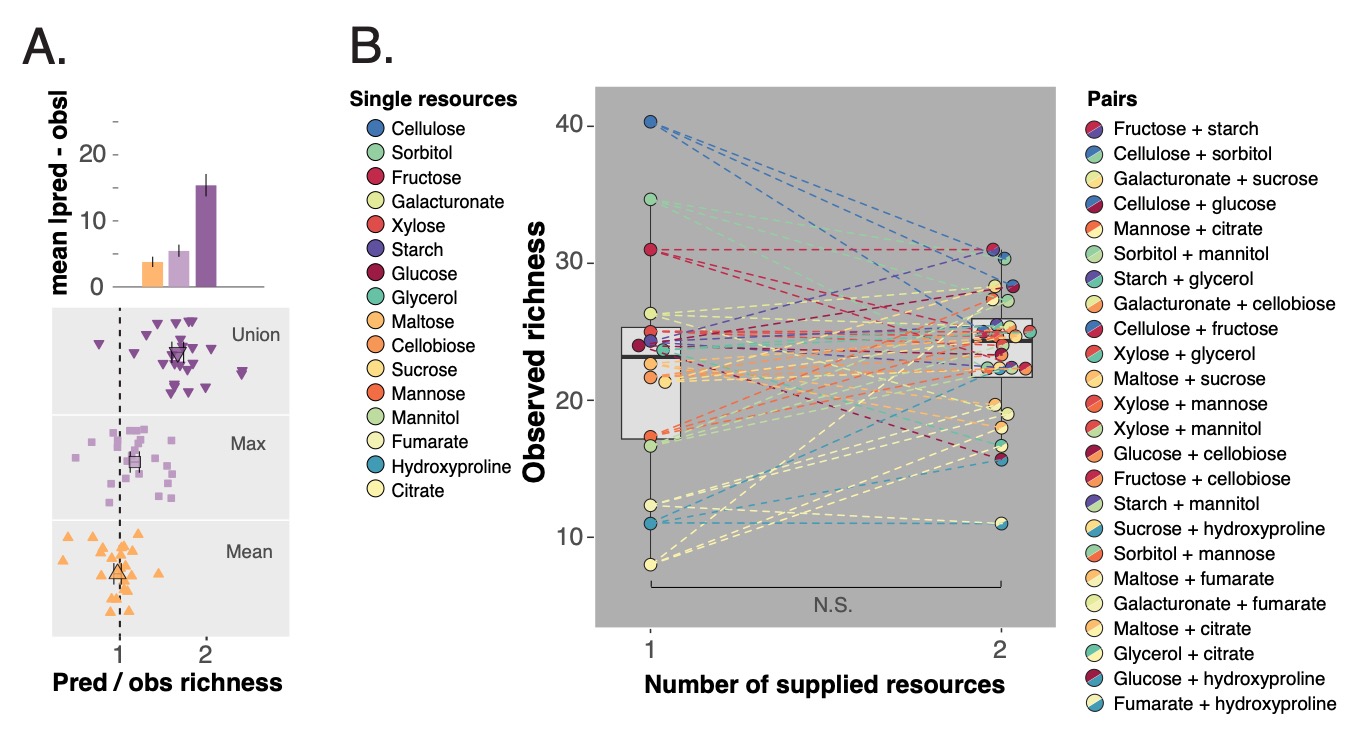

**Fig. S8. Richness of two-resource communities is approximately the average richness of constituent single-resource communities. A**. Observed richness of each two-resource community is best approximated by the average richness of constituent single resources, compared to the maximum and the union. Both the average error for the three predictions, calculated from the absolute values of predicted minus observed richness, and the ratio between predicted and observed richness are shown. **B**. Average richness of two-resource communities does not differ from the average richness of single-resource communities (boxplots with median and the 95 % confidence interval; number of 1-resource media = 16, number of 2-resource media = 24; each dot is obtained from the mean of 3 replicates, SEM are not shown for clarity). The color of the dots indicates the supporting resource(s).

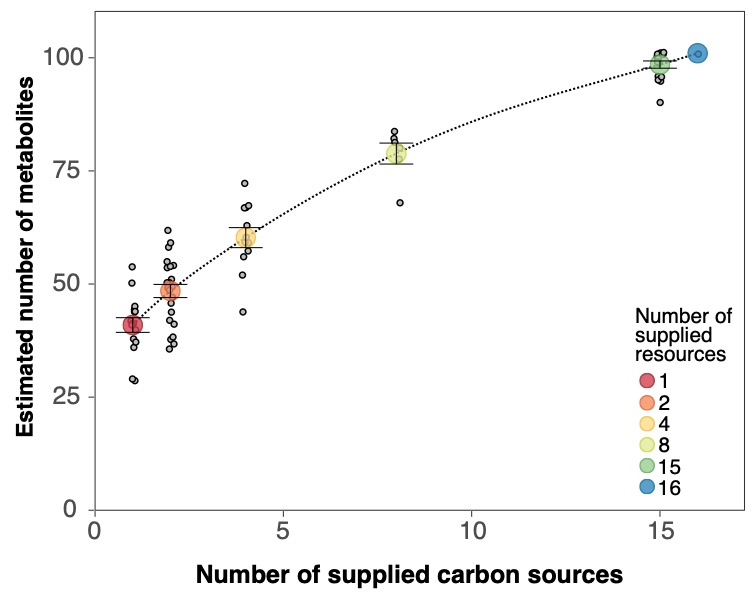

**Fig. S9.** **The estimated number of metabolites for combinations of carbon sources increases fast and tends to saturate with the number of supplied resources.** The number of metabolites has been computed from KEGG and MetaCyc databases (Materials and methods). Large colored dots indicate the average number of metabolite for each number of supplied resources (mean $\pm$ SEM) while small grey dots indicate the average richness in each media containing a combination of resources (16 for single-resource, 24 for two-resource, 12 for four-resource, six for eight-resource, 16 for 15-resource and one for 16-resource combinations). Error bars are omitted for clarity. The dotted line was obtained by fitting a spline to the points.

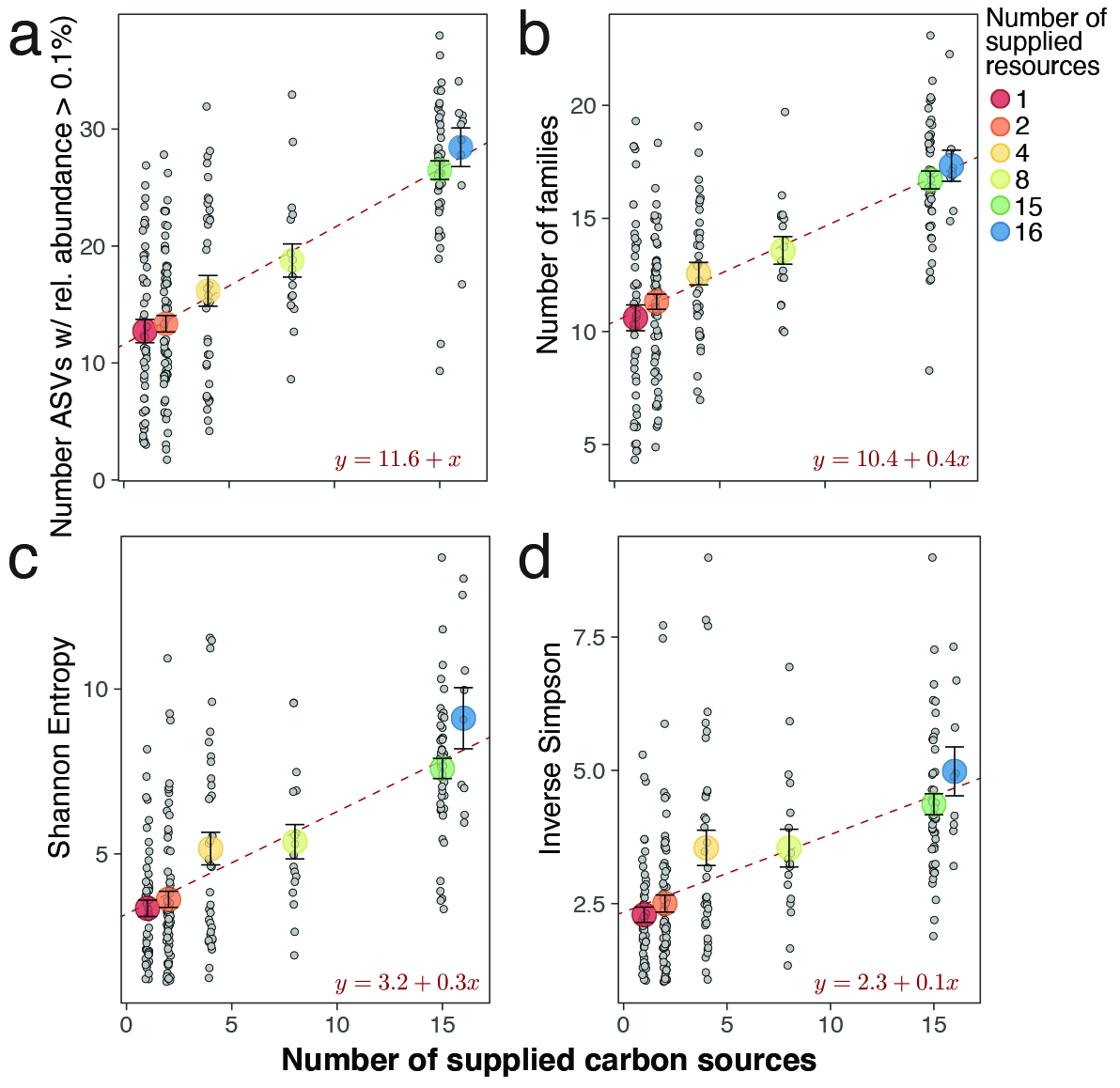

**Fig. S10. The observed linear trend is sufficiently robust to the exclusion of low-abundance ASVs, coarse-graining at the family level, and the index used to measure microbial community diversity.** **A.** Richness was calculated as the number of ASVs after the exclusion of those with relative abundance lower than 0.1%. **B.** Richness was calculated as the number of unique families in the media. **C, D.** The increase in diversity, measured as Shannon Entropy and Inverse Simpson Index, with the number of carbon sources can still be approximated by a line. These indices give progressively more weight to abundant species, accounting, in this way, for the evenness of the communities. In each panel, large colored dots indicate the mean $\pm$ SEM while small grey dots indicate the average richness in each media containing a combination of resources (16 for single-resource, 24 for two-resource, 12 for four-resource, six for eight-resource, 16 for 15-resource and one for 16-resource combinations). Error bars are omitted for clarity.

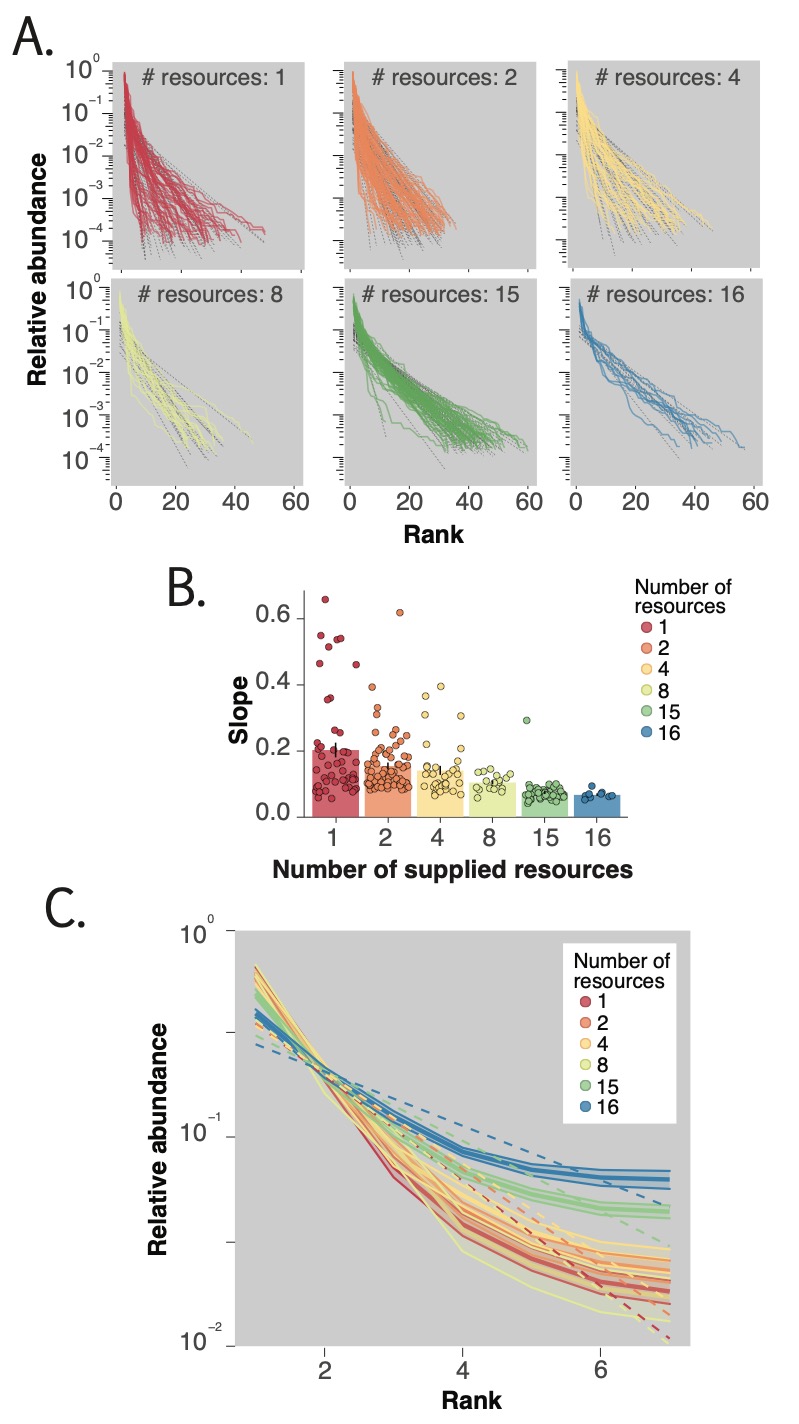

**Fig. S11. The evenness of the microbial communities increases with the number of supplied carbon sources.** **A**. Log-linear rank abundance distributions (RADs) are shown for all the experimental microbial microcosms (48 for single-resource, 72 for two-resource, 36 for four-resource, 18 for eight-resource, 16 for 15-resource and nine for 16-resource combinations) together with the fitted regression lines (black dashed lines). Going from one to 16 resources, RADs exhibit heavier tails. **B**. The average absolute value of the slope (bars indicate mean $\pm$ SEM across replicates with the same number of supplied resources, while jittered dots represent the slope for each individual replicate) decreases with the number of supplied resources. **C**. Changes in evenness are independent from changes in richness, as revealed by RADs normalized for richness (mean RADs, dashed colored lines, $\pm$ SD, shaded colored ribbon, for each number of supplied resources).

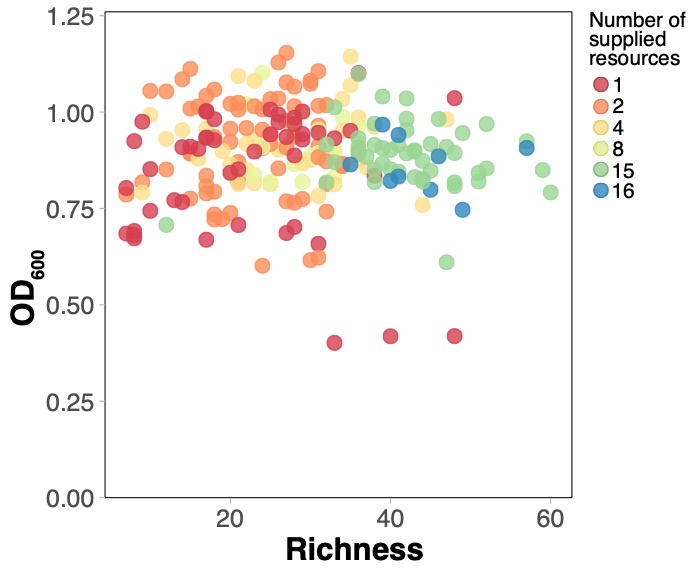

**Fig. S12. Biomass measured as OD_600_ does not change with community richness.** Colored dots represent OD values for each sample (48 for single-resource, 72 for two-resource, 36 for four-resource, 18 for eight-resource, 16 for 15-resource and nine for 16-resource combinations). The three points below 0.50 are the communities grown on cellulose.

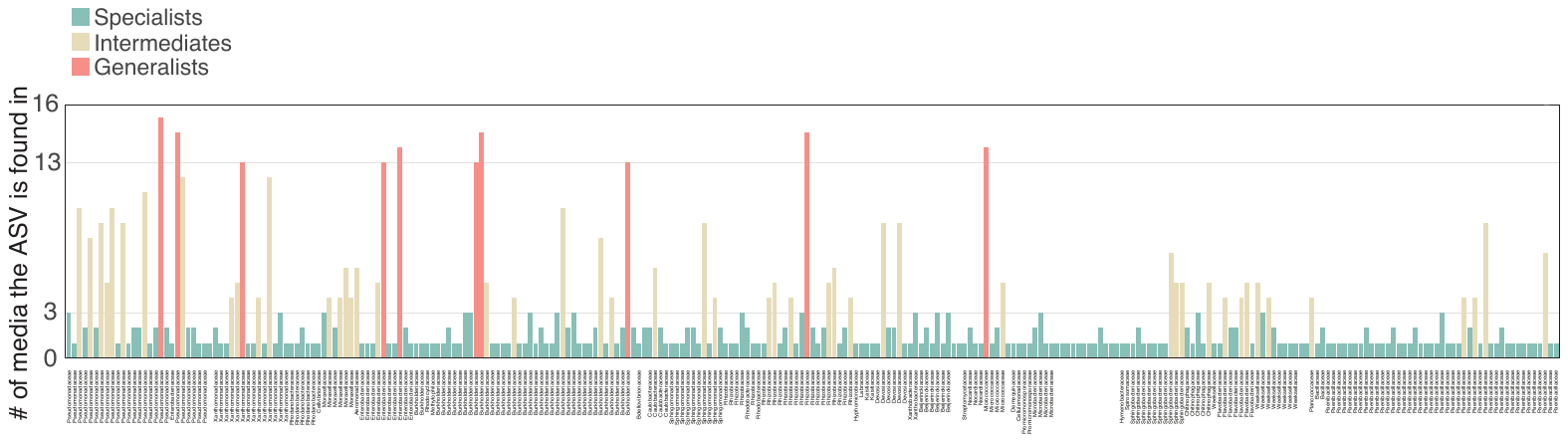

**Fig. S13. Resource occupancy of the 275 ASVs found in media containing a single carbon source.** The histogram shows the number of single resources in which each ASV is found. Bars are colored depending on whether the ASV has been classified as a generalist, a specialist or an intermediate. The families to which ASVs belong are reported.

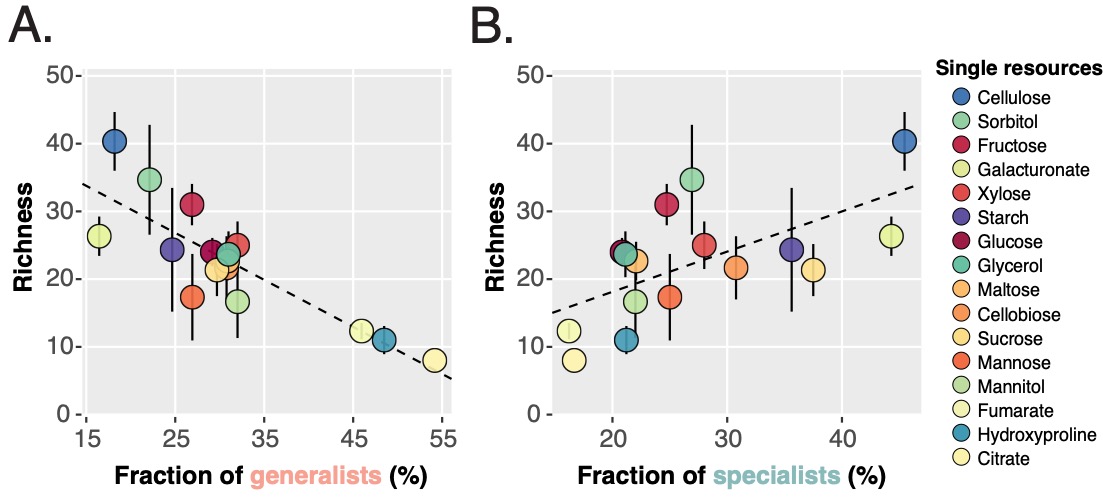

**Fig. S14. Fraction of generalist and specialist ASVs as a function of richness in single resources. A.** The fraction of generalist ASVs decreases with richness. **B.** The fraction of specialist ASVs in the community increases with richness in single resources. Fitted linear regression lines are shown.

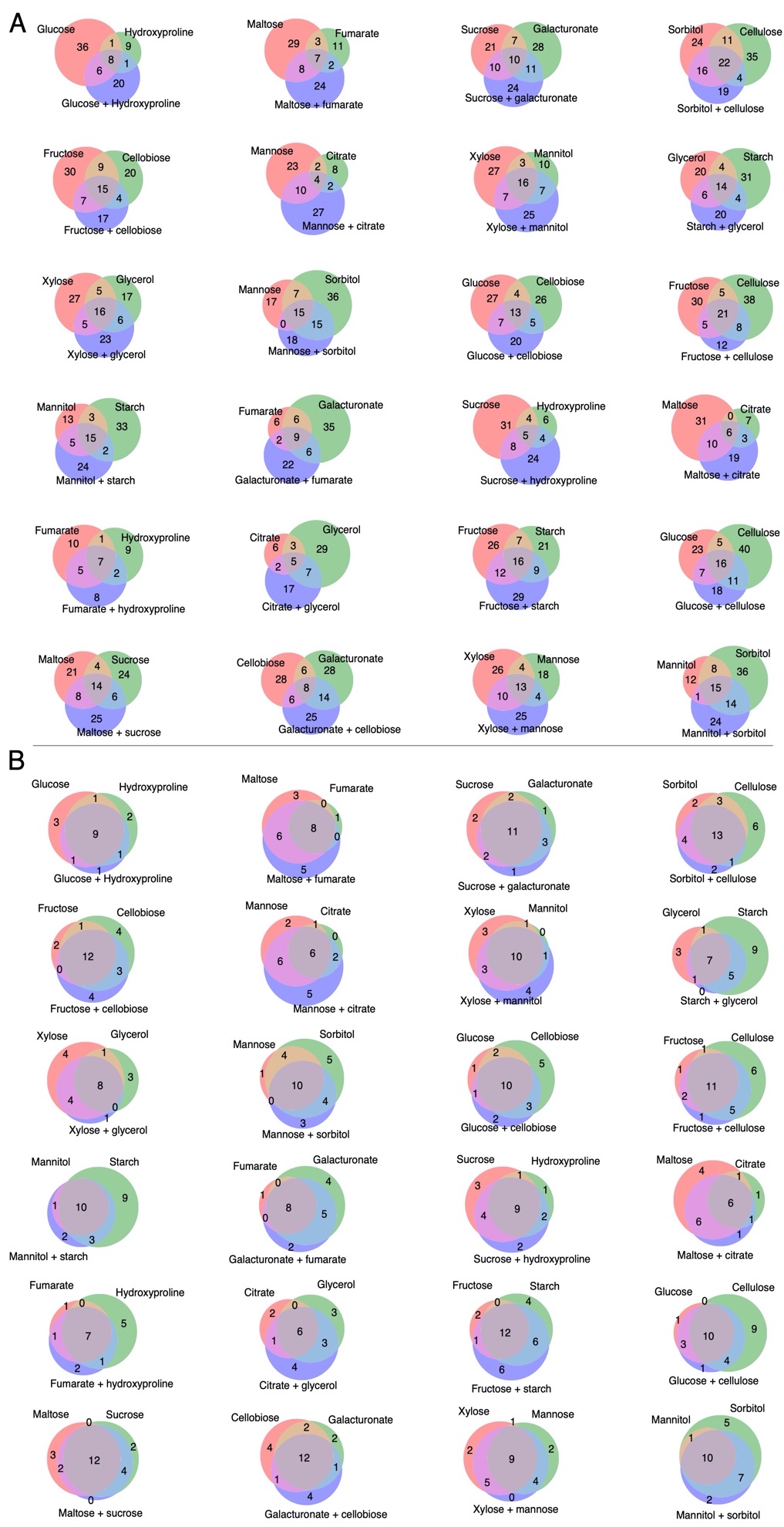

**Fig. S15. In the majority of two-resource communities, the number of ASVs not observed in constituent singles is larger than the number of ASVs shared by constituent singles. At the family level, instead, there is large overlap between constituent singles and correspondent two resource communities. A.** Venn diagrams of the union of replicates for each two-resource community are reported, showing the shared ASVs between the two constituents single-resource communities (light brown) and the ASVs not found in either of the constituent communities (blue). Red and green sections depict ASVs present only in one of the two constituents, while purple and light blue depict the ASVs share by one constituent community and the two-resource community. Note that red and green regions (ASVs that were not present in two-resource communities, can be large. **B**. The same with the number of families.

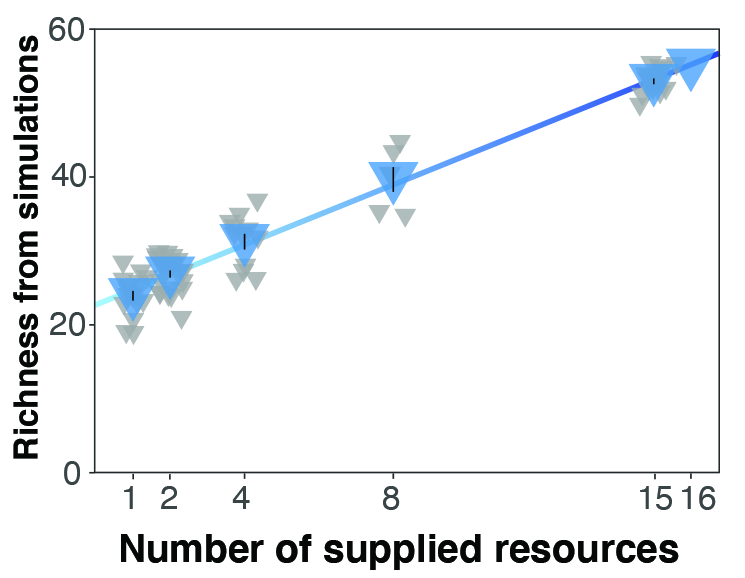

**Fig. S16. Richness from simulations grows modestly with the number of resources.** Richness obtained from simulations of the consumer-resource model with cross-feeding (blue triangles, mean $\pm$ SEM, N = 16 for single-resource, 24 for two-resource, 12 for four-resource, six for eight-resource, 16 for 15-resource and 1 for 16-resource combinations) as a linear function of the number of available resources (solid blue line, intercept = 22.7, slope = 2). Grey jittered triangles indicate the richness of communities grown on a particular resource combination.

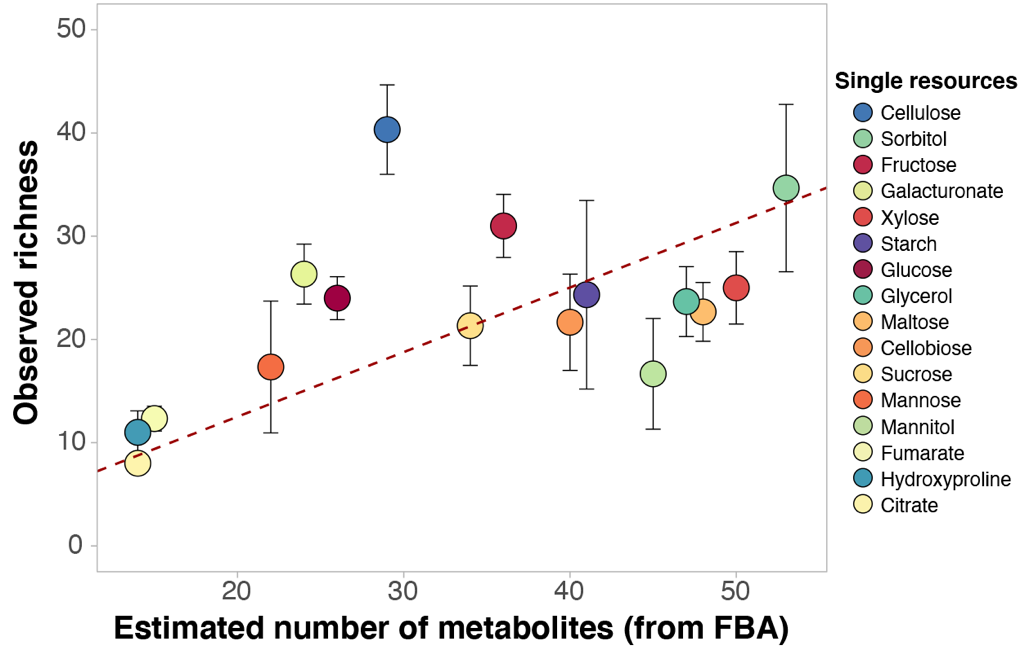

**Fig. S17. The number of metabolic byproducts predicted with FBA (flux balance analysis) correlates with community richness in single resources.** The estimated number of metabolic byproducts from single-resource communities, predicted using community-scale flux simulations (see Methods; Pearson correlation coefficient 0.52). Error bars indicate the SEM from the observed community richness data. The dashed line indicates the best-fit regression line.

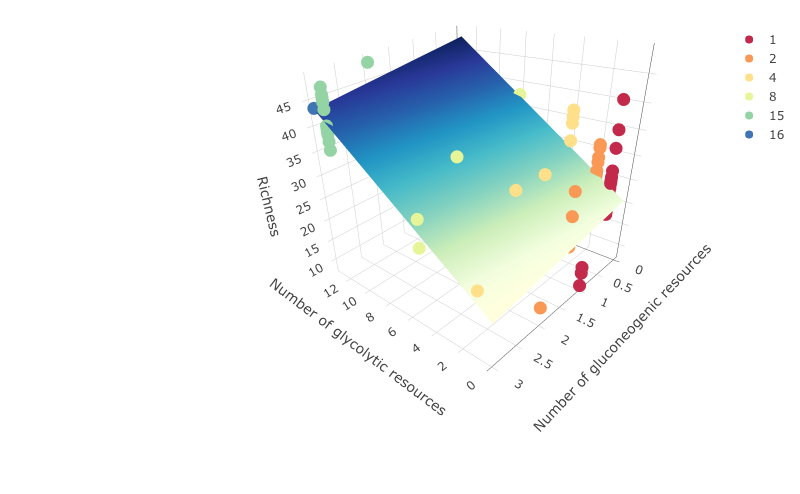

**Fig. S18. Glycolytic and gluconeogenic resources contribute differently to community richness.** The plot shows the relation between richness vs. the number of glycolytic (from 1 to 13 — glucose, fructose, xylose, mannose, cellobiose, maltose, sucrose, glycerol, galacturonate, cellulose and starch) and the number of gluconeogenic resources (from 1 to 3 — citrate, fumarate and hydroxyproline) in the growth media. While each glycolytic resource contributed ~ 2 species on top of the intercept (intercept = $20.73$, coefficient = $2.09 \pm0.21, p<0.001$), gluconeogenic resources did not increase community richness (coefficient = $- 1.49 \pm0.78, p<0.1$). Coefficient were estimated using a multivariate linear regression. Colors mark different total number of resources supplied in the growth media (from 1 to 16). The plane of predicted values is generated using the obtained regression coefficients.

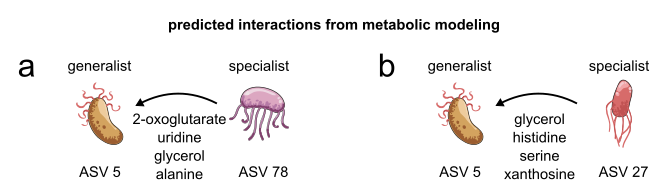

**Fig. S19. Habitat specialists are predicted to leak metabolic byproducts that can be used by habitat generalists.** Cartoons showing the predicted direction of metabolic interactions between generalist-specialist pairs found in our experimental communities. Predictions were made using pairwise flux-balance simulations (see Methods). Shown are two pairs as examples with metabolic interactions between **A.** ASV 5 (generalist) and ASV 78 (a specialist), and **B.** ASV 5 and ASV 27 (another specialist). The arrows show the direction of the predicted metabolic interactions, with the four exchanged metabolites with the highest cross-feeding potential marked below.

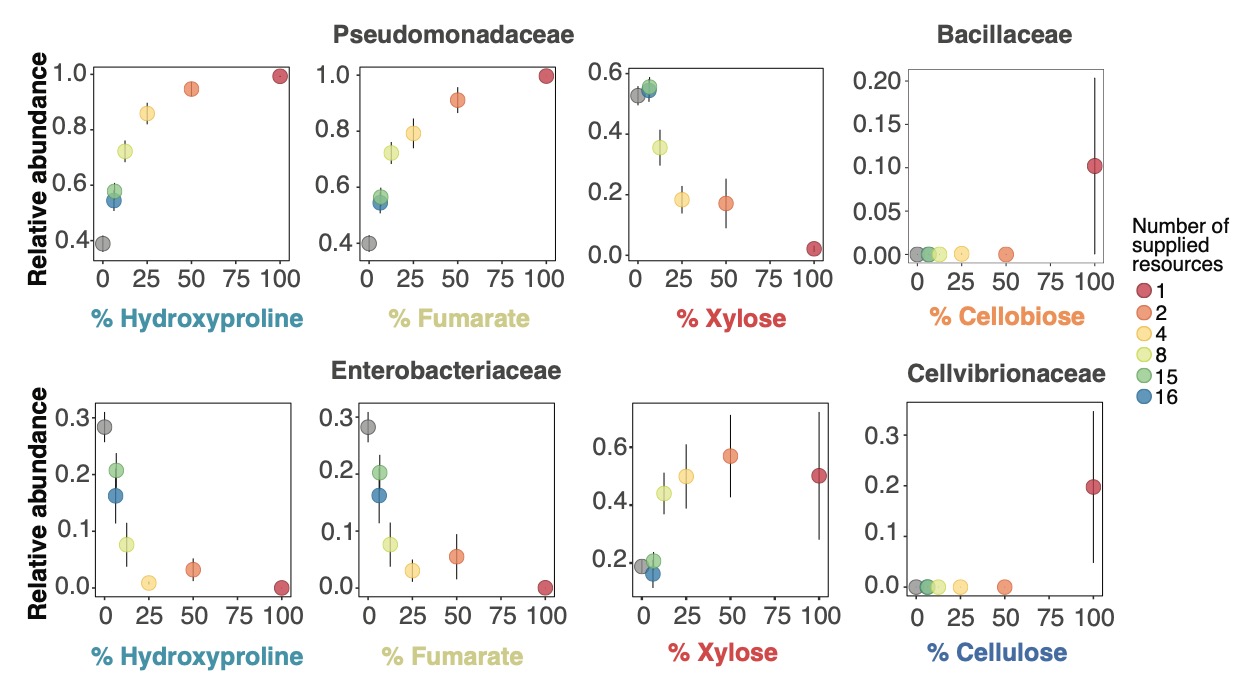

**Fig. S20. The abundance of the most prevalent bacterial families depends on the concentration of one or multiple resources of the pool supplied in the growth media.** The mean family abundance $\pm$ SEM for each relative concentration of the most relevant resource(s) identified through an ensemble tree regression (see Materials and methods) is shown. Families mostly composed of specialist taxa, e.g., Cellvibrionaceae and Bacillaceae, showed abrupt changes in their abundance with the concentration of the “favorite” resource. By contrast, more generalist families, e.g., Pseudomonadaceae and Enterobacteriaceae, exhibited smooth trends in their abundance with the concentration of multiple resources

**Table S1.** Detailed list of all the resource combinations used in the experiment.

| **Number of resources** | **Resource(s)** | **Relative concentration of each resource (%)** |
| --- | --- | --- |
| 1 | D-(+)-glucose (glucose) | 100 |
| 1 | D-(–)-fructose (fructose) | 100 |
| 1 | D-(+)-xylose (xylose) | 100 |
| 1 | D-(+)-mannose (mannose) | 100 |
| 1 | D-(+)-cellobiose (cellobiose) | 100 |
| 1 | D-(+)-maltose monohydrate (maltose) | 100 |
| 1 | sucrose | 100 |
| 1 | citric acid (citrate) | 100 |
| 1 | fumaric acid (fumarate) | 100 |
| 1 | D-(+)-galacturonic acid monohydrate (galacturonate) | 100 |
| 1 | D-mannitol (mannitol) | 100 |
| 1 | D-sorbitol (sorbitol) | 100 |
| 1 | glycerol | 100 |
| 1 | trans-4-Hydroxy-D-proline (hydroxyproline) | 100 |
| 1 | methyl cellulose (cellulose) | 100 |
| 1 | starch | 100 |
| 2 | glucose, hydroxyproline | 50 |
| 2 | maltose, fumarate | 50 |
| 2 | galacturonate, sucrose | 50 |
| 2 | sorbitol, cellulose | 50 |
| 2 | cellobiose, fructose | 50 |
| 2 | citrate, mannose | 50 |
| 2 | xylose, mannitol | 50 |
| 2 | glycerol, starch | 50 |
| 2 | glycerol, xylose | 50 |
| 2 | sorbitol, mannose | 50 |
| 2 | cellobiose, glucose | 50 |
| 2 | cellulose, fructose | 50 |
| 2 | starch, mannitol | 50 |
| 2 | fumarate, galacturonate | 50 |
| 2 | maltose, citrate | 50 |
| 2 | sucrose, hydroxyproline | 50 |
| 2 | hydroxyproline, fumarate | 50 |
| 2 | citrate, glycerol | 50 |
| 2 | starch, fructose | 50 |
| 2 | cellulose, glucose | 50 |
| 2 | sucrose, maltose | 50 |
| 2 | galacturonate, cellobiose | 50 |
| 2 | mannose, xylose | 50 |
| 2 | sorbitol, mannitol | 50 |
| 4 | glucose, hydroxyproline, maltose, fumarate | 25 |
| 4 | galacturonate, sucrose, sorbitol, cellulose | 25 |
| 4 | cellobiose, fructose, citrate, mannose | 25 |
| 4 | xylose, mannitol, glycerol, starch | 25 |
| 4 | glycerol, xylose, sorbitol, mannose | 25 |
| 4 | cellobiose, glucose, cellulose, fructose | 25 |
| 4 | starch, mannitol, fumarate, galacturonate | 25 |
| 4 | maltose, citrate, sucrose, hydroxyproline | 25 |
| 4 | hydroxyproline, fumarate, citrate, glycerol | 25 |
| 4 | starch, fructose, cellulose, glucose | 25 |
| 4 | sucrose, maltose, galacturonate, cellobiose | 25 |
| 4 | mannose, xylose, sorbitol, mannitol | 25 |
| 8 | glucose, hydroxyproline, maltose, fumarate, galacturonate, sucrose, sorbitol, cellulose | 12.5 |
| 8 | cellobiose, fructose, citrate, mannose, xylose, mannitol, glycerol, starch | 12.5 |
| 8 | glycerol, xylose, sorbitol, mannose, cellobiose, glucose, cellulose, fructose | 12.5 |
| 8 | starch, mannitol, fumarate, galacturonate, maltose, citrate, sucrose, hydroxyproline | 12.5 |
| 8 | hydroxyproline, fumarate, citrate, glycerol, starch, fructose, cellulose, glucose | 12.5 |
| 8 | sucrose, maltose, galacturonate, cellobiose, mannose, xylose, sorbitol, mannitol | 12.5 |
| 15 | all – glucose | 6.67 |
| 15 | all – fructose | 6.67 |
| 15 | all – xylose | 6.67 |
| 15 | all – mannose | 6.67 |
| 15 | all – cellobiose | 6.67 |
| 15 | all – maltose | 6.67 |
| 15 | all – sucrose | 6.67 |
| 15 | all – citrate | 6.67 |
| 15 | all – fumarate | 6.67 |
| 15 | all – galacturonate | 6.67 |
| 15 | all – mannitol | 6.67 |
| 15 | all – sorbitol | 6.67 |
| 15 | all – glycerol | 6.67 |
| 15 | all – hydroxyproline | 6.67 |
| 15 | all – cellulose | 6.67 |
| 15 | all - starch | 6.67 |
| 16 | all | 6.25 |
